# Thermal Proteomics and AI-assisted Target Deconvolution Identify ACLY as a Direct Target of Demethylzeylasteral in Psoriasis

**DOI:** 10.64898/2026.04.07.717093

**Authors:** Qiqi Wang, Nianzhou Yu, Yu Song, Xiaqiong Fan, Jing Tian, Qiong Yang, Yeye Guo, Chris Soon Heng Tan, Hongchao Ji

## Abstract

Target deconvolution of bioactive natural products (NPs) is frequently hampered by the inability of conventional thermal shift assays to distinguish direct ligand binding from indirect proteomic stabilization. Here, we developed an integrated target discovery framework that combines matrix-augmented pooling strategy-isothermal shift assay (MAPS-iTSA) thermal proteomics with a structure-aware graph neural network (HoloGNN). This strategy identified ATP-citrate lyase (ACLY) as a high-confidence target of Demethylzeylasteral (DEM), a triterpenoid from Tripterygium wilfordii. Orthogonal biochemical assays confirmed direct binding of DEM to ACLY (surface plasmon resonance, KD = 9.86 μM) and potent enzymatic inhibition (IC_50_ = 3.91 μM). To relate this interaction to the pharmacology of the source herb, thermal profiling of Tripterygium wilfordii extract combined with ACLY-based affinity-ultrafiltration mass spectrometry identified DEM as an ACLY-binding constituent of the extract. Given the established role of ACLY-mediated lipid metabolic reprogramming in psoriasis, we next evaluated the pharmacological significance of this interaction. DEM suppressed keratinocyte proliferation in vitro and, in vivo, alleviated imiquimod-induced psoriasiform dermatitis and reduced inflammatory cytokine expression. Single-cell RNA sequencing further revealed reversal of ACLY-SREBP-associated lipogenic reprogramming in keratinocytes. Untargeted lipidomics of lesional skin showed concordant reversal of the psoriatic lipid profile, including de novo synthesized fatty acids and acylcarnitines. Collectively, these findings establish ACLY as a direct and functionally relevant target of DEM and provide a robust AI-chemoproteomic paradigm for mechanism-guided NP drug discovery.

## INTRODUCTION

Natural products have long served as a rich source of bioactive compounds and therapeutic leads, offering unmatched structural diversity and biological potency^1^. Derived from a wide range of natural resources including plants, microbes, and marine organisms, these compounds have contributed significantly to modern therapeutics, particularly in oncology^2^. Classic examples such as paclitaxel and camptothecin highlight the potential of natural scaffolds to yield potent anti-cancer agents^3^. However, the intrinsic chemical complexity and polypharmacological behavior of these molecules often obscure the identification of their primary protein targets. This knowledge gap poses a major bottleneck to their clinical translation and rational development into optimized therapeutic agents^4^.

Derivatization-free chemoproteomic methods have emerged as powerful strategies to dissect the mechanisms of action of natural products without requiring compound immobilization or chemical modification. These include drug affinity responsive target stability (DARTS)^5^, limited proteolysis-mass spectrometry (LiP-MS)^6^, stability of proteins from rates of oxidation (SPROX)^7^, and thermal proteome profiling (TPP)^8–10^. Among these, TPP measures ligand-induced shifts in protein thermal stability across the proteome and has been widely applied. More recently, matrix-augmented pooling strategy-isothermal shift assay (MAPS-iTSA) has enabled simultaneous profiling of dozens of compounds in a single experiment, representing one of the highest-throughput derivatization-free platforms for large-scale target deconvolution of diverse natural product libraries^11^. Despite rapid expansion, thermal shift-based approaches are not without limitations. Because thermal stabilization reflects changes in protein stability rather than direct physical contact, proteins may display apparent stabilization due to indirect mechanisms, including downstream pathway perturbation, altered protein-protein interactions, metabolite redistribution or cellular stress responses^12^. Such effects can generate secondary or false-positive signals that do not arise from direct ligand binding, thereby complicating mechanistic interpretation^13^. Accordingly, orthogonal validation and computational frameworks that help distinguish direct from indirect stabilization events are increasingly essential to enhance confidence and mechanistic resolution in proteome-wide target discovery.

In parallel, the rise of artificial intelligence (AI) and geometric deep learning has transformed our ability to model molecular recognition^14^. A variety of sequence-, graph-, and structure-based models have been developed to predict protein-ligand interactions and binding affinities^15–21^. Among them, structure-aware models are particularly attractive because they can better capture spatial geometry and atom-level interaction patterns within protein-ligand complexes. However, these computational approaches have rarely been integrated with proteome-wide biophysical screening for target deconvolution of complex natural products. Such integration could improve hit prioritization and help resolve whether proteome-wide stabilization events reflect direct binding or indirect downstream effects.

Here, we developed an integrative experimental-computational framework that bridges thermal proteomics with structure-aware interaction modeling to systematically uncover and validate the targets of natural products. By combining MAPS-iTSA with a structure-aware graph neural network framework (HoloGNN), we established an integrated pipeline to refine, prioritize, and mechanistically interpret candidate protein-ligand interactions. We applied this approach to a panel of 50 structurally diverse natural products whose target spectra remain incompletely defined.

Among the top-ranked compound-target pairs from this integrated screen, the pair formed by Demethylzeylasteral (DEM), a triterpenoid derived from *Tripterygium wilfordii,* and the metabolic enzyme ATP-citrate lyase (ACLY) emerged as a robust hit. *Tripterygium wilfordii* extracts are known for their anti-inflammatory^22^ and anti-proliferative^23^ activities in conditions such as psoriasis, a disease in which ACLY-driven lipogenic reprogramming is implicated. ACLY is a central metabolic enzyme that catalyzes the ATP-dependent cleavage of citrate into oxaloacetate and acetyl-CoA, thereby linking carbohydrate metabolism to lipid biosynthesis, membrane production, and epigenetic regulation^24^. In addition to its established role in tumor metabolism^25^, accumulating evidence indicates that ACLY-driven metabolic reprogramming also contributes to inflammatory disorders, including psoriasis^26^, where aberrant lipogenesis is associated with keratinocyte hyperproliferation and inflammatory activation. Therefore, this discovery may reveal a new mechanism of action for *Tripterygium wilfordii* in treating psoriasis.

We therefore characterized this interaction mechanistically and pharmacologically. Biochemical assays confirmed that DEM directly binds and inhibits ACLY. Cellular studies showed that the compound depletes ACLY-derived acetyl-CoA, as read out by histone acetylation, and that its antiproliferative effect requires ACLY. In an imiquimod (IMQ)-induced mouse model of psoriasiform dermatitis, DEM alleviated skin inflammation and attenuated the ACLY-SREBP (sterol regulatory element-binding protein) lipogenic program in lesional skin. Together, these findings identify ACLY as a direct and functionally relevant target of DEM and suggest that the integration of MAPS-iTSA with structure-aware computational prioritization provides a generalizable strategy for mechanism-driven discovery from bioactive natural products.

## RESULTS

### An integrated experimental–computational workflow for target prioritization

In this study, we established a streamlined experimental-computational workflow integrating MAPS-iTSA screening, HoloGNN-assisted prioritization, and downstream biochemical confirmation to identify candidate direct targets of natural products (**Fig.1**). Structurally diverse natural products were distributed into overlapping pools according to a predefined sensing matrix. Following thermal treatment and mass spectrometry-based protein quantification, pool-resolved abundance profiles were deconvoluted to nominate candidate compound-protein pairs. These candidates were then assessed using docking-derived complex structures and HoloGNN, which combined local interaction features with attention-based structural representations to generate predicted binding scores. This structure-aware step provided complementary evidence for prioritizing thermally responsive proteins, without equating thermal stabilization with direct binding. Selected interactions were subsequently subjected to surface plasmon resonance and drug affinity responsive target stability assays for orthogonal confirmation. Together, these stages linked multiplexed proteomic screening with structural assessment and biochemical validation, providing a systematic route from natural product libraries to experimentally supported target assignments.

**Fig. 1.**
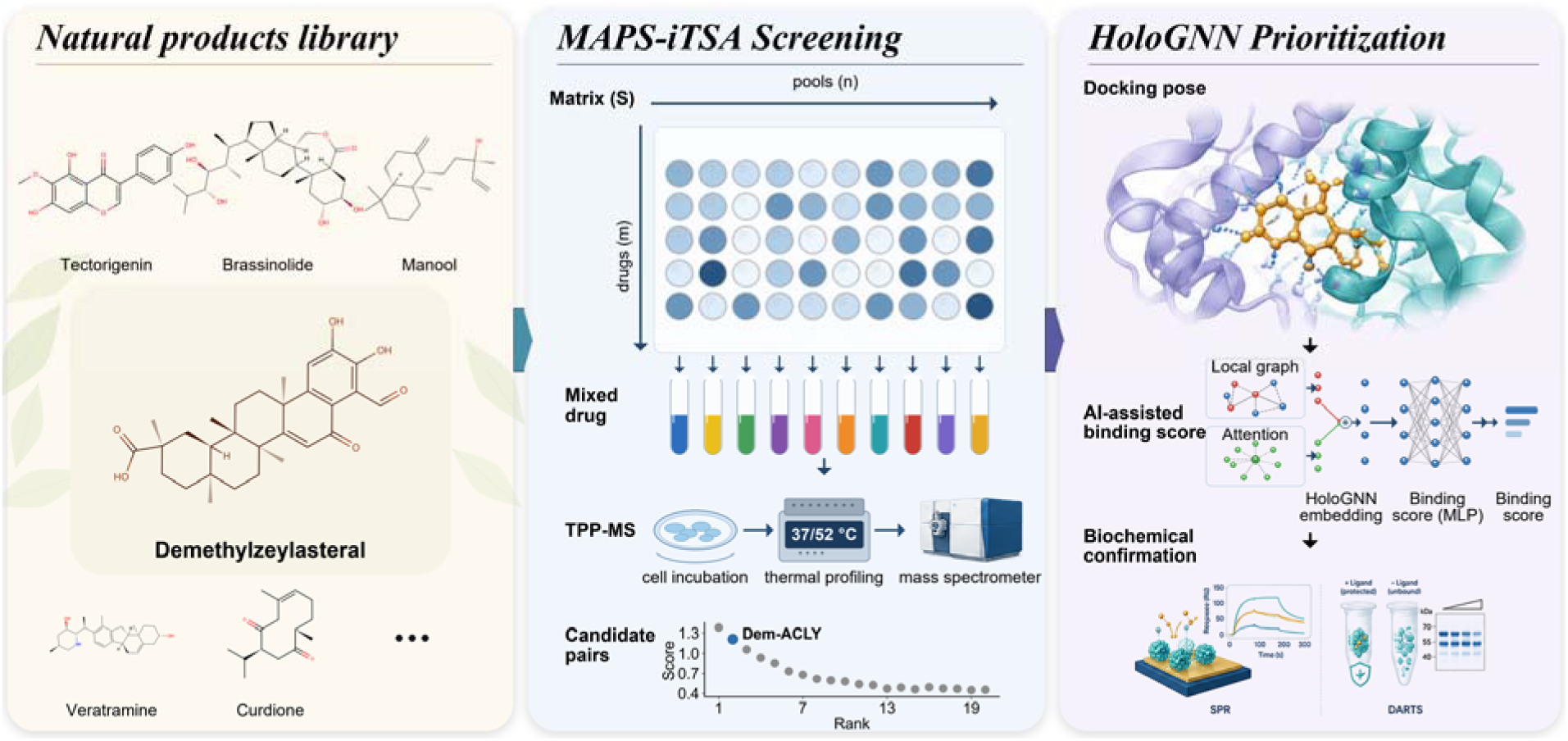
An integrated experimental-computational framework for target deconvolution of natural products. (1) A library of structurally diverse natural products. (2) MAPS-iTSA: compounds are pooled according to a sensing matrix, lysates are thermally challenged, the soluble proteome is quantified by LC-MS/MS, and pool-resolved profiles are deconvoluted to rank candidate compound-protein pairs. (3) HoloGNN scores the docked complexes to prioritize pairs for biochemical confirmation by SPR and DARTS.

### ACLY is identified as a target of Demethylzeylasteral

Using a panel of 50 structurally diverse natural products (**Table S1**), MAPS-iTSA generated candidate compound-target pairs from pooled isothermal proteome profiling. Screening was performed in K562 cell lysates, which provide a high-yield, well-characterized human proteome with the broad coverage required for matrix-pooled thermal profiling, rather than in intact cells, with all 50 compounds tested at an identical final concentration of 20 μM (1% DMSO) and pooled five per tube according to a 15 × 25 matrix design, so that each compound occupied a unique combination of pools. In each experiment, approximately 4,000 proteins were identified (**Table S2 and S3**). **Fig.2a** illustrates the quantitative correlation among the 15 pools, with Pearson correlation coefficients consistently exceeding 0.98. Because non-target proteins are expected to have similar abundances across pools, the high correlations indicate that the experimental procedure was highly reproducible and stable.

**Fig. 2.**
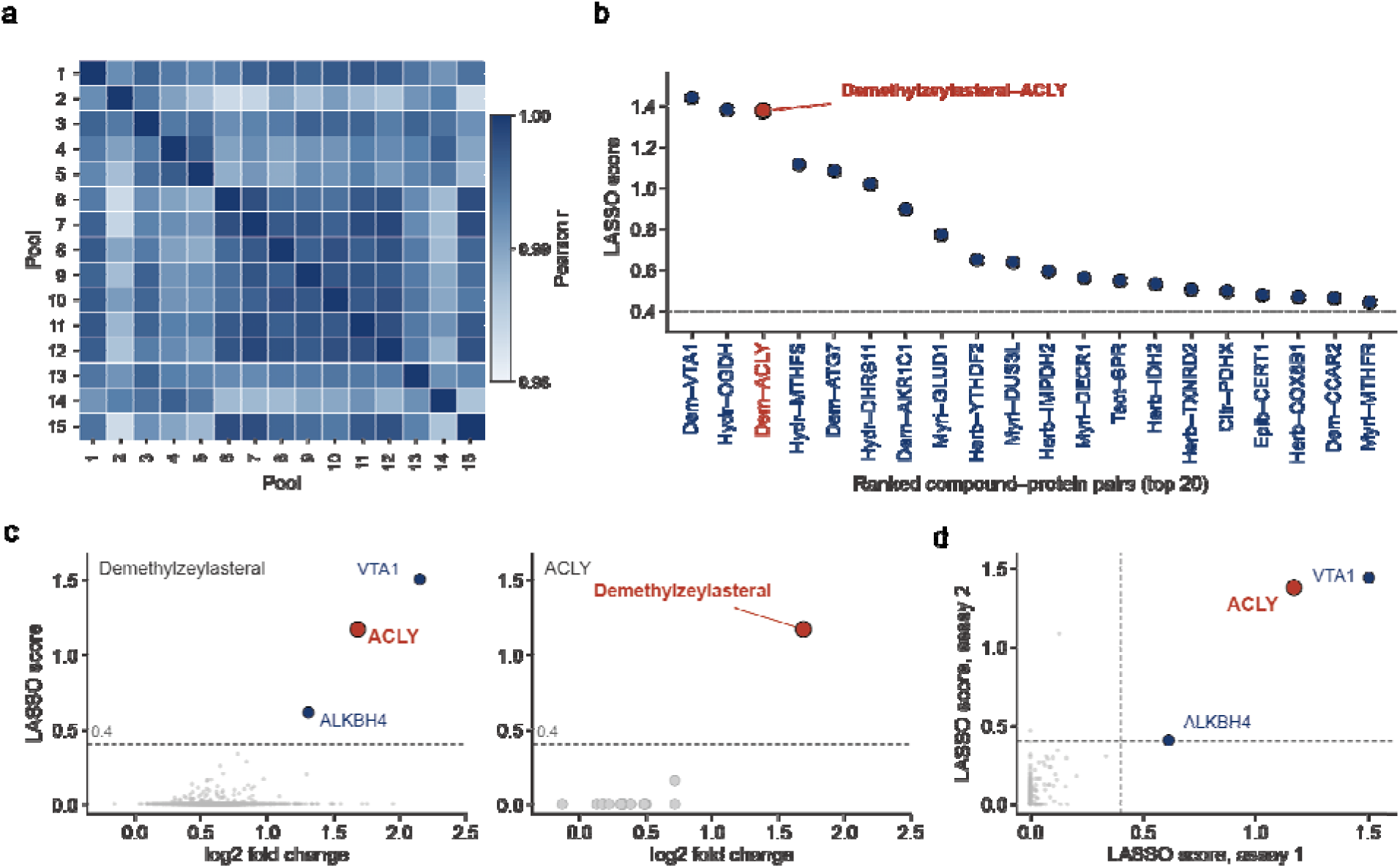
Identification of ACLY as a candidate target of Demethylzeylasteral. **a**. Heatmap reflects the reproducibility across the pooled samples of MAPS-iTSA. **b**. Drug-target confidence scores derived from MAPS-iTSA experiments calculated by LASSO algorithm. **c**. Relationship between log2 fold change and LASSO scores for identified targets. **d**. Reproducibility of drug-target confidence scores across replicate experiments.

We next summarized the top-ranking natural product-target pairs from the first MAPS-iTSA screen, in which the DEM-ACLY pair emerged among the high confidence candidates (**Fig.2b**). On the basis of this convergent workflow, ACLY was prioritized for subsequent biochemical and functional validation. To further assess the specificity of this interaction, we plotted log2 fold change against the interaction score for DEM across all identified proteins, where ACLY was positioned among the top-ranking candidates with both a high interaction score and a pronounced thermal-shift signal (**Fig.2c**, left). Conversely, when ACLY was evaluated across the screened natural products, DEM ranked at the top and showed clear enrichment relative to the remaining compounds (**Fig.2c**, right).

In addition, comparison of interaction scores between the two independent experiments, which were performed on separate days using lysates prepared from two independently thawed and cultured batches of K562 cells, demonstrated strong reproducibility of the deconvoluted hits, with ACLY remaining a robust candidate across replicates (**Fig.2d, Table S4**). These reciprocal observations provide converging evidence that the DEM-ACLY pair is highly likely to represent a true biological interaction, thereby justifying the subsequent focus on ACLY in downstream validation experiments.

### HoloGNN prioritization supports direct binding of Demethylzeylasteral to ACLY

To further determine whether the DEM-ACLY pair identified by MAPS-iTSA was compatible with a direct physical interaction, we next performed structure-aware computational prioritization using HoloGNN. Docking analysis placed DEM in a plausible ACLY binding pocket, where the compound formed multiple close-contact interactions with surrounding residues, including GLN-385, ASN-383, and ARG-379, supporting a structurally feasible binding configuration (**Fig.3a**).

**Fig. 3.**
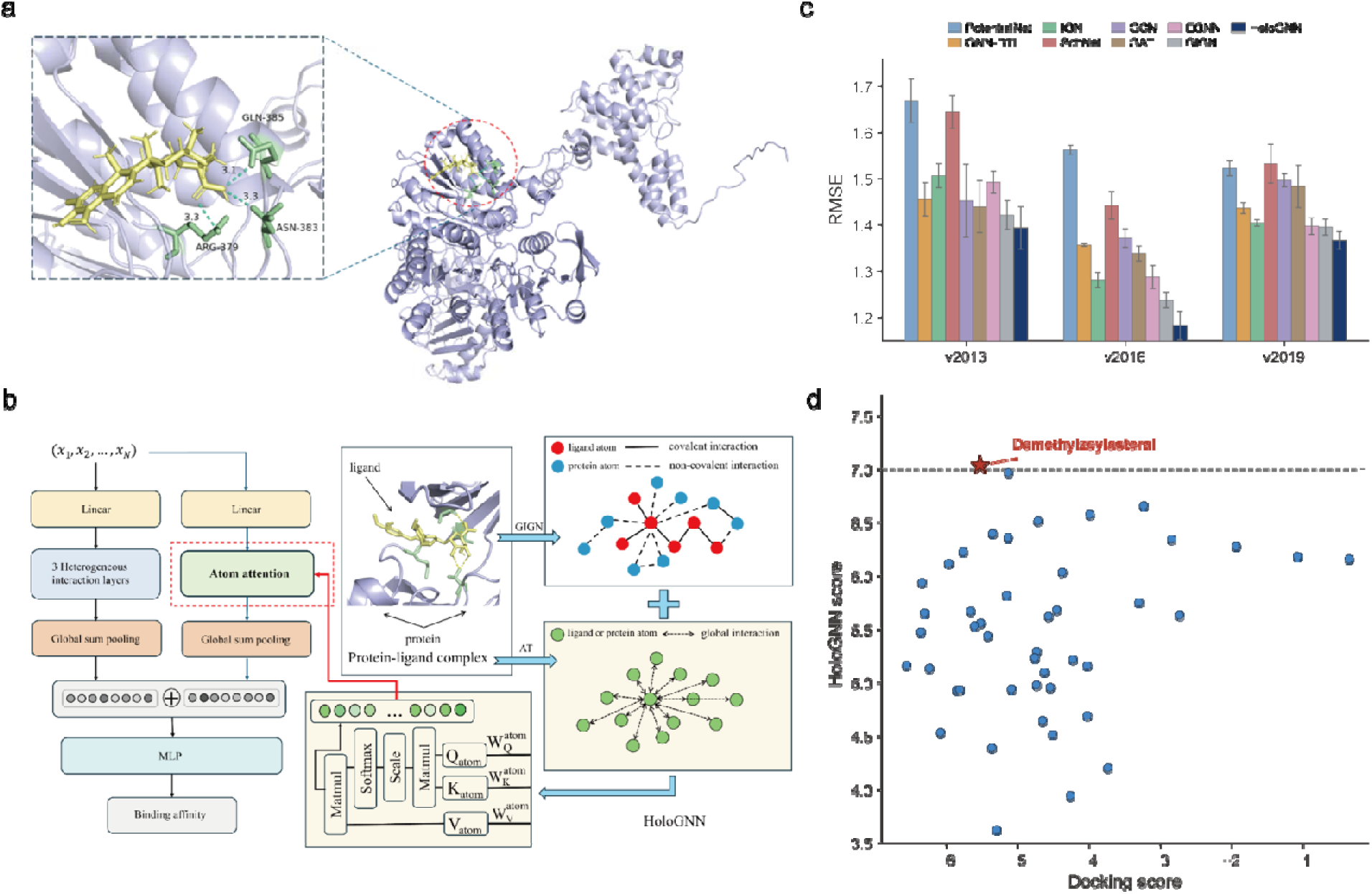
HoloGNN prediction and docking supports direct Demethylzeylasteral-ACLY binding. **a.** Representative docking pose of the DEM-ACLY complex. **b**. Brief schematic of HoloGNN model architecture. **c**. RMSE values comparison between HoloGNN and SOTA (state-of-the-art) models. **d**. Scatter plot of docking scores versus HoloGNN-predicted binding affinities for 50 natural products targeting ACLY.

HoloGNN represents the protein-ligand complex as a unified atomic graph that jointly captures local interaction patterns and global structural context. Local interaction graph features and holographic attention features are integrated through feature fusion and subsequently processed by a regressor to generate a binding score, enabling prioritization of compound-target pairs with a higher likelihood of direct binding (**Fig.3b**). To evaluate model performance, we benchmarked HoloGNN against representative methods across three PDBbind test sets, where it consistently achieved the lowest RMSE (**Fig.3c**) and highest Pearson correlation (**Fig.S1a**) values among the evaluated models, supporting its robustness for structure-based binding prediction. To further evaluate HoloGNN in a biologically relevant setting, we assessed its ability to predict recently reported drug-target interactions, where it accurately prioritized known ligands for BRAF, CDK11, and CSNK2A2 (**Fig.S1b-d**)^11,27^.

We then applied this framework to the 50 natural products included in our screen and predicted their binding scores against ACLY using docking-derived complex structures as input (**Table S5**). As shown in **Fig.3d**, DEM was positioned among the top-scoring compounds in the joint distribution of docking score and HoloGNN score, indicating concordance between favorable structural accommodation and a high model-predicted binding propensity. Notably, the DEM-ACLY pair that emerged from the MAPS-iTSA screen was also prioritized by HoloGNN, providing orthogonal computational support for this interaction and suggesting that the observed thermal stabilization is more likely to reflect direct target engagement rather than an indirect downstream effect. Together, these results support the inference that DEM directly engages ACLY and provide a rationale for subsequent biochemical validation of this candidate interaction.

### Biochemical analyses confirm Demethylzeylasteral inhibits ACLY

To independently validate the predicted DEM-ACLY interaction and determine whether ACLY represents a direct biochemical target of DEM, we performed a series of orthogonal biochemical assays with DEM tested as a single agent rather than within a compound pool. Cellular thermal shift assay (CETSA) performed in the cell-lysate format showed that DEM treatment stabilized endogenous ACLY in HaCaT cell lysates, with significantly elevated protein retention at 52°C (P < 0.01) and 55°C (P < 0.001) compared with vehicle controls, indicating target engagement under thermal challenge (**Fig.4a**).

**Fig. 4.**
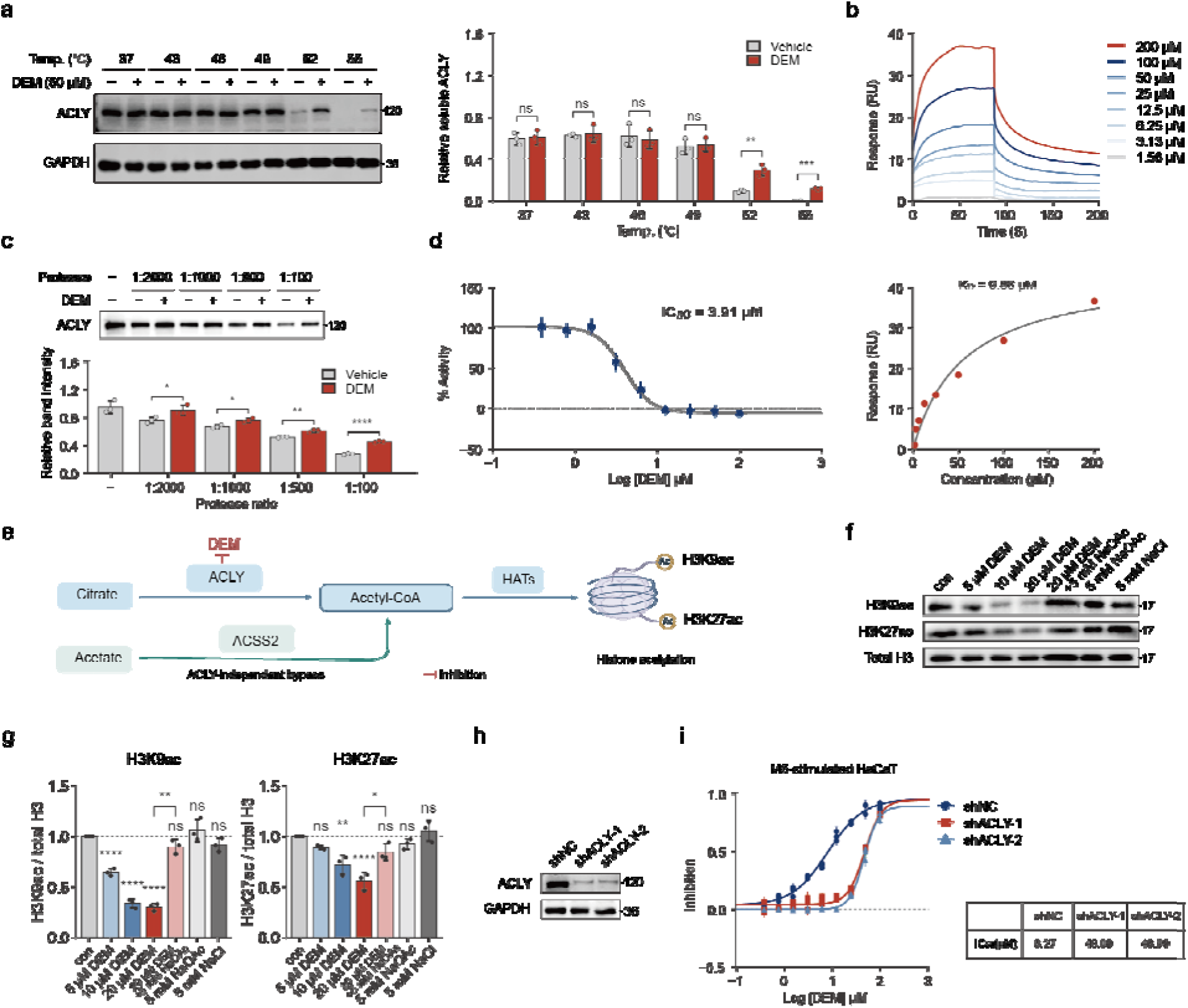
Biochemical validation of the Demethylzeylasteral-ACLY interaction. **a.** CETSA (cell-lysate format) of endogenous ACLY in HaCaT cell lysates incubated with DEM (50 μM). Left, representative blots; right, quantification. **b.** SPR analysis of DEM binding to immobilized ACLY. Top, sensorgrams (1.56-200 μM); bottom, steady-state affinity fit (K_D_ = 9.86 μM). **c.** DARTS assay of ACLY protection against pronase digestion. Top, representative blots; bottom, quantification. **d.** Inhibition of ACLY enzymatic activity (IC = 3.91 μM, four-parameter logistic fit). **e.** The ACLY-acetyl-CoA-histone acetylation axis and the ACSS2-dependent acetate bypass tested in f and g. **f.** Representative blots of H3K9ac, H3K27ac and total histone H3 in HaCaT keratinocytes treated for 24 h as indicated. **g.** Densitometric quantification of f, normalized to total histone H3 in the same lane and to the vehicle control of the same experiment (vehicle = 1). **h.** Immunoblot confirming ACLY knockdown in HaCaT cells stably expressing shNC, shACLY-1 or shACLY-2. i. DEM dose-response curves in M5-stimulated HaCaT cells expressing shNC, shACLY-1 or shACLY-2 (72 h); IC values are listed beside the curves. Inhibition was normalized to the vehicle control of the corresponding cell line. Data are mean ± s.e.m. of three independent biological replicates, with individual replicates shown as symbols. Significance: two-tailed unpaired Student’s *t*-test (a, c); one-way ANOVA with Dunnett’s test against vehicle plus a pre-specified paired comparison of DEM versus DEM plus acetate (bracket) (g). ns, not significant; *P < 0.05, **P < 0.01, ***P < 0.001, ****P < 0.0001.

To further determine whether DEM interacts with ACLY through direct physical binding, we next employed surface plasmon resonance (SPR) and drug affinity responsive target stability (DARTS) assays. SPR analysis revealed concentration-dependent binding responses across a 1.56-200 μM range, yielding a dissociation constant (K_D_) of 9.86 μM (**Fig.4b**), thereby demonstrating direct binding of DEM to purified ACLY. Consistent with this result, DARTS assays showed that DEM conferred dose-dependent protection of ACLY against pronase digestion at all protease-to-protein mass ratios tested (1:2000 to 1:100; P < 0.05 to P < 0.0001), further supporting ligand-induced conformational stabilization of ACLY upon compound binding (**Fig.4c**).

Having established direct binding, we next assessed whether this interaction also produced a functional consequence at the enzymatic level. DEM inhibited ACLY activity in a dose-dependent manner, with an IC_50_ of 3.91 μM (**Fig.4d**), indicating that binding of the compound is accompanied by efficient suppression of ACLY catalytic activity. Notably, the SPR-derived K_D_ (9.86 μM) and the enzymatic IC_50_ (3.91 μM) fall within the same low-micromolar range, reinforcing the conclusion that DEM not only physically engages ACLY but also exerts a direct functional inhibitory effect.

Because ACLY is the principal source of nucleocytosolic acetyl-CoA, the obligate acetyl donor for histone acetyltransferases, cellular ACLY catalytic function can be read out directly from histone acetylation, which can be restored independently of ACLY by exogenous acetate through ACSS2 (**Fig.4e**). Using this relationship, we next asked whether DEM inhibits ACLY within intact cells. Treatment of HaCaT keratinocytes for 24 h reduced histone acetylation in a dose-dependent manner (**Fig.4f-g**). H3K9ac declined significantly at all three concentrations (P < 0.0001 for each), whereas H3K27ac reached significance only from 10 μM onwards (P = 0.0014 and P < 0.0001 at 10 and 20 μM; **Fig.4g**). Co-treatment with 5 mM sodium acetate, which regenerates acetyl-CoA through ACSS2 and thereby bypasses ACLY, restored histone acetylation in DEM-treated cells, recovering 85% of the H3K9ac deficit (P = 0.002) and 65% of the H3K27ac deficit (P = 0.018), such that neither mark differed significantly from vehicle after rescue (P = 0.21 and P = 0.08, respectively). Sodium acetate alone and an equimolar NaCl osmotic control left histone acetylation unchanged relative to vehicle (P > 0.3 for all), excluding a contribution of acetate itself, of sodium or of osmolarity to this rescue. Together these data indicate that the decline in histone acetylation reflects depletion of ACLY-derived acetyl-CoA.

We then asked whether the antiproliferative activity of DEM likewise depends on ACLY. HaCaT cells stably expressing either of two independent ACLY-targeting shRNAs (shACLY-1 and shACLY-2; approximately 90% knockdown at the protein level; **Fig.4h**) or a non-targeting control shRNA (shNC) were compared in a psoriasis-like model induced by the M5 cytokine cocktail (TNF-α, IL-17A, IL-22, IL-1α and oncostatin M). ACLY knockdown shifted the dose-response curve rightward in both knockdown lines, increasing the IC from 8.27 μM in shNC cells to 48.09 μM in shACLY-1 cells and 46.99 μM in shACLY-2 cells, corresponding to a 5.7-5.8-fold loss of potency (**Fig.4i**).

Taken together, these convergent biochemical and cellular data demonstrate that DEM directly engages and binds ACLY, stabilizing the protein against both thermal and proteolytic challenge. The accompanying loss of catalytic activity, the dose-dependent decline in histone acetylation that is rescued by exogenous acetate, and the ACLY-dependence of the antiproliferative effect in keratinocytes together establish ACLY as a direct and functionally relevant molecular target of DEM.

Beyond this single pair, we asked whether the same screen supports the recovery and validation of additional interactions. Pool-to-pool reproducibility in the second compound set was comparable to that of the first **(Fig.S2a)**, and the Chrysosplenol D-ATIC pair was among the highest-scoring compound-protein pairs after LASSO deconvolution **(Fig.S2b)**: ATIC was the top-scoring protein for Chrysosplenol D and Chrysosplenol D the only compound above threshold for ATIC **(Fig.S2c, d),** and ATIC scored highest in both independent screens **(Fig.S2e).** Orthogonal assays, performed in an unrelated cellular background (AC16 cells) and with purified protein, confirmed the interaction. DARTS showed significant protection of ATIC against pronase digestion at protease-to-protein mass ratios of 1:1000 and 1:500 (P < 0.01), with protection no longer detectable at the highest protease concentration tested (1:100) **(Fig.S2f)**. SPR gave a dissociation constant of 21.86 μM **(Fig.S2g)**, and CETSA in the cell-lysate format showed significant stabilization of endogenous ATIC from 52 °C onwards (P < 0.01 at 52, 55 and 58 °C) **(Fig.S2h)**. Recovery and biochemical confirmation of a second, mechanistically unrelated pair indicates that the MAPS-iTSA screening and deconvolution workflow is not specific to the DEM-ACLY interaction.

### Demethylzeylasteral-ACLY engagement links to the anti-psoriatic activity of *Tripterygium wilfordii*

*Tripterygium wilfordii* has been used in the treatment of psoriasis, and both clinical evidence and experimental studies support its anti-psoriatic activity^28^. To determine whether the DEM-ACLY interaction identified above is relevant in the context of the source herb, we next examined *Tripterygium wilfordii* extract using complementary target-engagement approaches.

Isothermal shift assay (iTSA) of the extract identified ACLY as an extract-responsive protein showing positive thermal stabilization, indicating that ACLY is not only engaged by the purified compound but is also detectable at the herbal extract level. Although ACLY did not rank among the most significant hits at the extract level, it exhibited reproducible positive stabilization and was accompanied by two convergent signals, FASN and VTA1, both of which were highlighted in the extract-level analysis (**Fig.5a, Table S6, Table S7**). Among these, FASN is a key downstream lipogenic enzyme functionally linked to the ACLY-centered metabolic axis, whereas VTA1 was also identified as a candidate hit of DEM in the MAPS-iTSA dataset, thereby providing an additional point of overlap between the extract-level and monomer-level target landscapes. Notably, extract treatment significantly increased the relative abundance of ACLY, FASN, and VTA1, with VTA1 showing the most pronounced response, further supporting coordinated engagement of this ACLY-associated signaling context by the herbal extract (**Fig.5b**). To assess reproducibility across cellular backgrounds, we further compared the extract-level TPP responses in two independent cell lines. ACLY, FASN, and VTA1 showed broadly concordant significance patterns in THP-1 and HepG2 cells, indicating that these extract-responsive signals were reproducible across distinct cellular contexts rather than being restricted to a single cell line (**Fig.5c**).

**Fig. 5.**
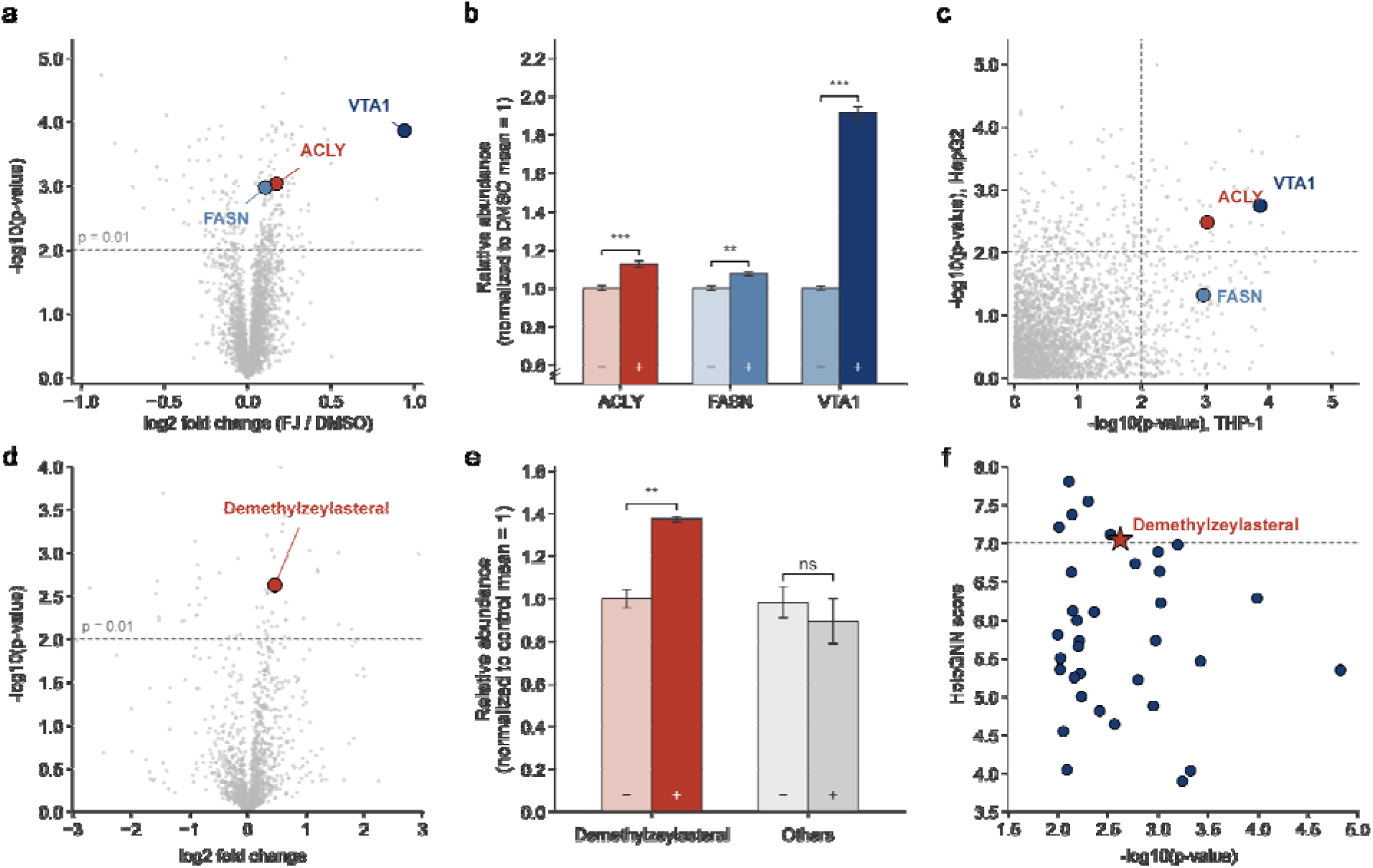
Demethylzeylasteral-ACLY engagement is supported in the context of *Tripterygium wilfordii* extract. **a.** Volcano plot of iTSA results from *Tripterygium wilfordii* extract, highlighting ACLY, FASN, and VTA1 as extract-responsive proteins; **b.** Relative abundance of ACLY, FASN, and VTA1 in the extract-treated group compared with DMSO control; **c.** Comparison of extract-level iTSA results in THP-1 and HepG2 cells. **d.** Volcano plot of ACLY-based AUMS showing selective enrichment of DEM in the ACLY-binding fraction; **e.** Relative abundance of DEM and other detected compounds in the ACLY-binding fraction; **f.** HoloGNN-based prioritization of AUMS-positive compounds, with DEM ranked as a high-scoring candidate among ACLY-binding components.

To further test whether DEM represents an ACLY-interacting constituent within the herbal context, we performed an ACLY-based affinity-ultrafiltration mass spectrometry (AUMS) assay^29^. In the AUMS dataset, DEM was significantly enriched in the ACLY-binding fraction, providing direct analytical support that DEM is one of the ACLY-binding components associated with *Tripterygium wilfordii* (**Fig.5d**). By contrast, the pooled group of other detected compounds did not show significant enrichment, further supporting the preferential association of DEM with ACLY within the extract-derived component set (**Fig.5e**). Consistent with this result, when AUMS-positive compounds were further evaluated by HoloGNN, DEM emerged as a prominent candidate with comparatively high predicted binding score, further prioritizing it among the extract-derived ACLY-binding components (**Fig.5f**).

### Demethylzeylasteral alleviates IMQ-induced psoriasis-like dermatitis in vivo

Given that the mechanism of ACLY in relation to tumors has been extensively studied^30,31^, we selected psoriasis as the disease model for pharmacological validation. ACLY catalyzes the conversion of citrate to acetyl-CoA and serves as the rate-limiting node for de novo fatty acid and cholesterol biosynthesis. Previous studies have demonstrated that ACLY is overexpressed in psoriatic lesions and that its inhibition attenuates keratinocyte proliferation and inflammatory responses; moreover, Acly-knockout mice exhibit defective skin barrier lipids, implicating ACLY as a key driver of lipid metabolic reprogramming in psoriasis^32,33^. These lines of evidence directly link ACLY to the aberrant lipid metabolism characteristic of psoriasis and suggest that pharmacological targeting of ACLY holds therapeutic potential. Given that DEM was shown to directly bind and inhibit ACLY enzymatic activity, we therefore employed psoriasis as the disease model to evaluate its pharmacological efficacy.

To evaluate the in vivo efficacy of DEM, female C57BL/6 mice were subjected to the IMQ-induced psoriasiform model, receiving daily topical 5% IMQ cream alongside concurrent intraperitoneal DEM (1 mg/kg) (Fig.6a). Compared with the IMQ group, mice in the IMQ+Demethylzeylasteral (IMQ+DEM) group exhibited markedly reduced erythema, scaling, and skin thickening, and Psoriasis Area and Severity Index (PASI) scores confirmed a significant decrease in disease severity (**Fig.6b, d**). At the molecular level, expression of *Ccl3*, *Il6*, and *Tnf* in lesional skin was significantly downregulated in the IMQ+DEM group relative to the IMQ group, indicating that DEM effectively suppressed the pro-inflammatory cytokine milieu that drives psoriatic pathology (**Fig.6e**). Importantly, while repeated IMQ application characteristically induces weight loss secondary to reduced food intake, DEM did not exacerbate this decline **(Fig.6c)**. Furthermore, histological evaluation by hematoxylin and eosin (H&E) staining revealed no morphological abnormalities across major organs (heart, liver, spleen, lung, and kidney), which, together with the stable body weight trajectory, confirms the highly favorable systemic safety profile of DEM **(Fig.6f)**.

**Fig. 6.**
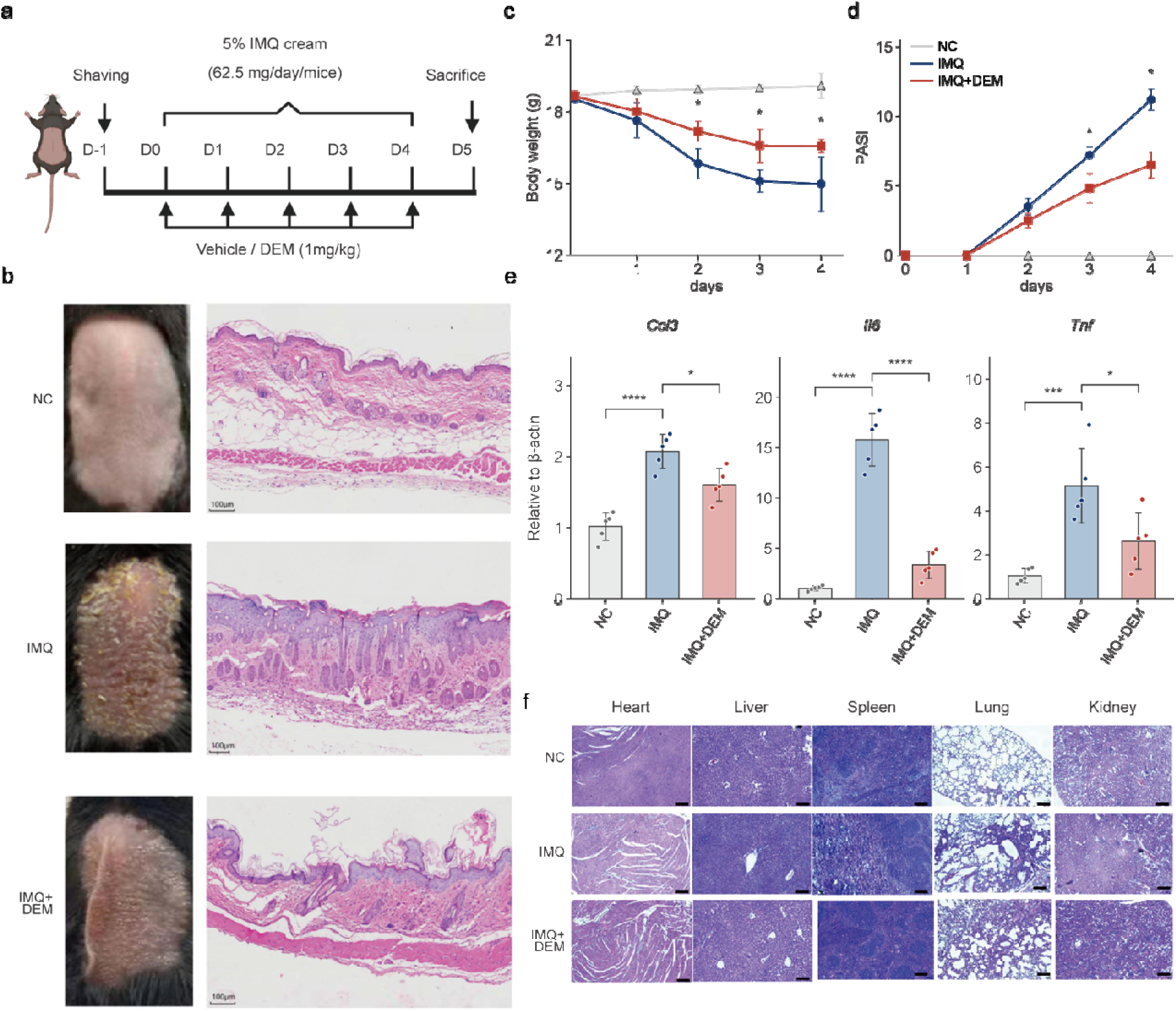
Demethylzeylasteral alleviates IMQ-induced psoriasis-like dermatitis. **a.** Experimental design for the IMQ-induced psoriasiform mouse model with concurrent DEM administration (1 mg/kg, i.p., daily). **b.** Representative macroscopic images of dorsal skin (left) and corresponding H&E-stained skin sections (right) across the NC, IMQ, and IMQ+DEM groups. Scale bars, 100 μm. **c-d.** Daily changes in body weight (c) and cumulative Psoriasis Area and Severity Index (PASI) scores (d) over the 4-day treatment period. **e.** Relative mRNA levels of *Ccl3*, *Il6*, and *Tnf* in lesional skin quantified by qRT-PCR. f. Representative H&E-stained sections of major organs (heart, liver, spleen, lung, and kidney) from each group, confirming no treatment-related systemic histopathological changes. Scale bars, 200 μm. Data in (c-e) are presented as mean ± SD (n = 5 biologically independent mice per group). Statistical significance was determined by two-tailed Student’s t-test. *P < 0.05, ***P < 0.001, ****P < 0.0001 versus the IMQ group. NC, normal control; IMQ, imiquimod; DEM, Demethylzeylasteral.

### Demethylzeylasteral reverses ACLY-dependent lipogenic reprogramming in psoriasiform keratinocytes

To dissect the mechanism of action of DEM at single-cell resolution, we performed scRNA-seq on dorsal skin from NC, IMQ, and IMQ+DEM mice. After stringent quality control, 10,639 (NC), 24,224 (IMQ), and 9,881 (IMQ+DEM) cells were retained for downstream analysis (**Fig.7a**). Uniform manifold approximation and projection (UMAP) embedding and cell proportion analysis revealed marked shifts in cellular composition across conditions, most notably within the keratinocyte (KC) compartment (**Fig.7a, b**). A dot plot of canonical lineage markers confirmed robust cell-type annotation: keratinocytes were enriched for Krt-family genes (Krt14, Krt10, Krt1), fibroblasts expressed Col1a1 and Dcn, endothelial cells were marked by Pecam1 and Cdh5, and myeloid cells showed high expression of Lyz2, among others (**Fig.S3c**). Further subclustering of keratinocytes revealed that IMQ induced a pronounced expansion of hyperproliferative (Cycling KC) and pro-inflammatory KC subsets, a shift that was substantially reversed by DEM treatment, restoring the KC subpopulation distribution toward that of normal skin (**Fig.S3a, b**).

**Fig. 7.**
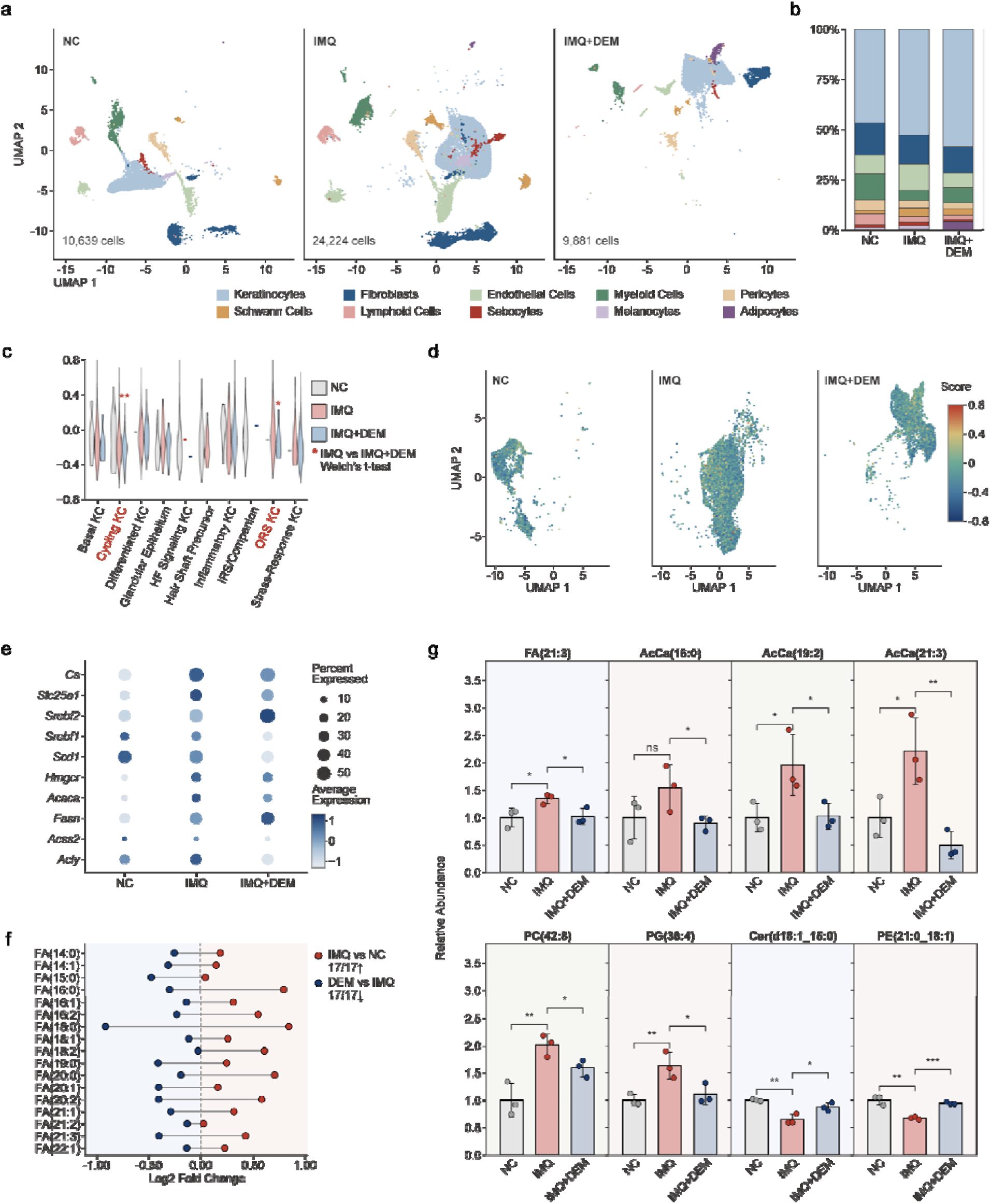
Demethylzeylasteral suppresses IMQ-induced ACLY pathway activation in psoriasiform keratinocytes at single-cell resolution. a-b. UMAP visualization (a) and cell proportion analysis (b) of skin cells from the NC, IMQ, and IMQ+DEM groups. **c.** ACLY pathway activity scores across different keratinocyte (KC) subtypes. **d.** UMAP density plots illustrating KC transcriptomic states across conditions. **e.** Dot plot showing the expression of ACLY pathway and lipogenesis-related genes in KCs. **f-g.** Untargeted lipidomics reveals the bidirectional shifts of 17 canonical de novo synthesized and elongated free fatty acids (f) and the relative abundance of representative ACLY-downstream lipid effectors (g). Data in (g) are presented as mean ± SD (n = 3). Significance: Welch’s t-test comparing IMQ with IMQ+DEM (c) and two-tailed unpaired Student’s t-test (g). *P < 0.05, **P < 0.01, ***P < 0.001; ns, not significant.

Gene set enrichment analysis (GSEA) performed on the ranked differentially expressed gene list of keratinocytes demonstrated significant positive enrichment of lipid metabolism-associated pathways (“Response to lipid”, P = 0.0009, adjusted P = 0.035), with the enrichment peak occurring at the leading edge of the ranked list, indicating activation of a lipogenesis-centered transcriptional program in IMQ-stimulated keratinocytes **(Fig.S3d)**. To resolve these metabolic changes at single-cell resolution, we evaluated ACLY pathway module scores across KC subtypes. While pathway activity varied across individual KC subsets, DEM treatment significantly reduced ACLY pathway scores in Cycling KC and ORS KC relative to IMQ **(Fig.7c)**. Consistently, feature-weighted UMAP projections showed that NC keratinocytes formed a compact, low-scoring cluster, whereas IMQ-treated keratinocytes exhibited a broadly dispersed distribution; conversely, the IMQ+DEM group contracted into a spatially restricted state predominantly characterized by low pathway scores **(Fig.7d)**. Profiling of ACLY-associated genes revealed a consistent transcriptional remodeling across treatment groups **(Fig.7e)**. While IMQ stimulation elevated core lipogenic and metabolic enzymes (*Acly*, *Cs*, *Acaca*, and *Hmgcr*), DEM treatment effectively suppressed these targets back to baseline levels, alongside sustained repression of *Srebf1*, *Scd1*, and *Acss2*. Together, these findings demonstrate that DEM restricts upstream metabolic supply and core enzymes, reshaping the ACLY-associated transcriptional program.

Because mRNA expression changes do not necessarily equate to alterations in metabolite abundance, we next performed untargeted LC-MS/MS lipidomics on lesional skin (n = 3 per group) in both positive and negative ionization modes to test whether DEM reverses ACLY-driven lipid metabolic reprogramming at the metabolic level. Across 1,911 quantified lipid species spanning 26 lipid classes, IMQ induced extensive lipidomic perturbation (335 differential lipids, P < 0.05). DEM treatment counter-regulated 109 species (|FC| > 1.2 in opposite directions, P < 0.05), suppressing 54 IMQ-upregulated lipids while restoring 55 IMQ-depleted lipids (**Fig.S4c-e**). Principal component analysis (PCA) of the DEM-responsive lipidome confirmed a clear separation between the NC and IMQ groups along PC1 (57.4% of variance), whereas the IMQ+DEM group shifted markedly back toward the NC cluster (**Fig.S4a**). Mechanistically, ACLY-derived cytosolic acetyl-CoA and malonyl-CoA serve as essential precursors for FASN-mediated de novo fatty acid synthesis and subsequent fatty acyl chain elongation. Strikingly, across all 17 canonical de novo synthesized and elongated free fatty acids (C14-C22, including palmitate FA(16:0), stearate FA(18:0), oleate FA(18:1), and the elongated species FA(20:0)-FA(22:1)), 100% (17/17) were elevated in IMQ lesions relative to NC controls, and 100% (17/17) were coordinately downregulated by DEM (**Fig.7f**). Similarly, acylcarnitines (AcCa)—which reflect mitochondrial acyl-CoA handling—exhibited a global accumulation in the IMQ group (39/44 species elevated, 88.6%), which was broadly reversed by DEM (37/44 species reduced, 84.1%; **Fig.S4b**). Specifically, DEM significantly attenuated downstream lipid effectors, reducing free fatty acids [FA(21:3)], palmitoylcarnitine [AcCa(16:0)], long/very-long-chain acylcarnitines [AcCa(19:2), AcCa(21:3), AcCa(24:4), AcCa(24:5)], and membrane phospholipids [PC(42:8), PC(44:7), PC(26:1_18:2), PG(36:4)] (P < 0.05; **Fig.7g, Fig.S4d**). Concurrently, DEMrescued the IMQ-induced depletion of essential epidermal barrier and structural lipids, including homeostatic ceramides [Cer(d18:1_15:0), Cer(d33:1)], sphingomyelin [SM(d43:1)], and phosphatidylethanolamines [PE(21:0_18:1), PE(25:0_18:1)] (P < 0.05; **Fig.7g, Fig.S4d, e**). Collectively, these integrated single-cell transcriptomic and untargeted lipidomic findings demonstrate that DEM suppresses the ACLY-SREBP lipogenic program and reverses pathological lipid metabolic remodeling in vivo, providing a direct metabolic basis for its amelioration of epidermal hyperproliferation and inflammation.

## DISCUSSION

In this study, we established an integrated target deconvolution strategy that combines MAPS-iTSA screening, HoloGNN-assisted structural prioritization, and orthogonal biochemical validation to identify direct targets of bioactive natural products. Using this workflow, we prioritized ACLY as a candidate target of DEM and subsequently demonstrated direct DEM-ACLY engagement through CETSA, SPR, DARTS, and enzymatic inhibition assays. Coupled with single-cell transcriptomics and untargeted lipidomics in an in vivo psoriasiform model, these findings define ACLY as a direct and functionally relevant target of DEM and illustrate how complementary proteomic, multi-omic, and structure-aware approaches can improve confidence in natural product target identification.

A central challenge in thermal proteomics-based target discovery is that protein stabilization does not necessarily indicate direct ligand binding. Our study addressed this limitation by introducing a prioritization step that integrates docking-based structural assessment and HoloGNN scoring with MAPS-iTSA-derived candidate pairs. Rather than relying on thermal shifts alone, this combined strategy enabled us to distinguish a high-confidence direct-binding candidate from broader proteome-wide stabilization events. In this context, HoloGNN functioned not as a stand-alone discovery engine, but as a structure-informed filter that increased confidence in selecting ACLY for downstream validation.

The identification of ACLY as a direct target of DEM is biologically and pharmacologically notable. ACLY catalyzes the ATP-dependent cleavage of citrate into oxaloacetate and acetyl-CoA in the cytosol, occupying a critical metabolic branchpoint that bridges central carbon metabolism with lipid biosynthesis and acetyl-CoA homeostasis^34^. Dysregulation of ACLY has been implicated in a broad spectrum of pathological conditions: in cancer, ACLY-derived acetyl-CoA fuels de novo lipogenesis and supports histone acetylation that drives oncogenic transcriptional programs^35,36^; in neurodegenerative disorders, altered ACLY-dependent metabolism contributes to neuronal dysfunction^37^; and in inflammatory skin diseases, ACLY-dependent metabolic reprogramming drives keratinocyte hyperproliferation and immune activation^33^. Although bempedoic acid, a prodrug activated predominantly in hepatocytes, has been approved for hypercholesterolemia^38,39^, its tissue-restricted activation limits extrahepatic utility, and no natural product-derived ACLY inhibitor has been reported to date.

DEM is a friedelane-type triterpenoid from *Tripterygium wilfordii* with documented anti-inflammatory and anti-proliferative activities, yet its direct molecular targets have remained incompletely defined^22^. Recent studies have begun to reveal its polypharmacological nature: in tumor immunology, DEM directly binds the deubiquitinase USP22 at Arg371, promoting USP22 degradation and PD-L1 ubiquitin-proteasome-dependent turnover to activate antitumor immunity^40^; in hepatocellular carcinoma, DEM has been reported to suppress liver cancer stem cell maintenance in association with reduced glycolysis-linked histone lactylation (H3K9la and H3K56la), although a direct protein target was not identified in that study^41^; and in lung cancer, DEM has been reported to bind LRPPRC and to inhibit the biogenesis of oxidative phosphorylation complexes^42^. Our thermal shift, enzymatic inhibition, and quantitative proteomic data now establish that DEM directly engages ACLY and suppresses its catalytic function, providing a mechanistic basis for its lipid metabolism-related bioactivities and expanding its known target repertoire, which includes the deubiquitinase USP22 and LRPPRC, to include a central enzyme of lipid metabolism. The reported targets of DEM differ in the level of validation achieved, ranging from a mapped binding residue (USP22) to phenotypic associations for which no direct protein target was assigned.

Establishing direct target engagement required a multi-tiered biochemical validation strategy. CETSA confirmed that DEM thermally stabilizes endogenous ACLY in HaCaT cell lysates, yet this assay alone cannot exclude indirect mechanisms. We therefore employed SPR with purified recombinant ACLY, which demonstrated direct, concentration-dependent binding (KD = 9.86 μM), and DARTS assays, which showed dose-dependent proteolytic protection consistent with ligand-induced conformational change. The IC_50_ for enzymatic inhibition (3.91 μM) falls within the same low-micromolar range as the SPR-derived KD, supporting a model in which the observed loss of catalytic activity arises from direct target binding. The convergence of these orthogonal approaches-target engagement in cell lysates (thermal shift), direct binding (SPR), conformational stabilization (DARTS), and functional inhibition (enzymatic assay)-collectively strengthens the evidence that ACLY is a direct and functionally relevant target of DEM. Inhibition of ACLY function within intact cells was established separately, through the dose-dependent loss of histone H3K9 and H3K27 acetylation that was rescued by exogenous acetate but not by an equimolar osmotic control, and through the rightward shift of the DEM dose-response curve in two independent ACLY-knockdown keratinocyte lines. Importantly, the additional evidence from *Tripterygium wilfordii* extract thermal profiling and ACLY-based AUMS links the DEM-ACLY interaction to the pharmacological context of the source herb, supporting ACLY engagement as one plausible contributor to its anti-psoriatic effects.

Psoriasis was selected as the validation model based on converging evidence positioning ACLY as a pathogenic driver. Psoriatic lesions exhibit pronounced lipid metabolic reprogramming with upregulation of ACLY-dependent biosynthetic pathways, and ACLY inhibition attenuates keratinocyte proliferation and inflammatory responses in vitro^43^. Despite this rationale, no ACLY-targeting therapeutic has entered clinical development for psoriasis. Our in vivo experiments demonstrated that DEM significantly attenuated IMQ-induced psoriasiform dermatitis, reducing PASI scores and suppressing *Ccl3*, *Il6*, and *Tnf* expression without detectable organ toxicity, consistent with the proposed mechanism of ACLY inhibition.

Integrating single-cell RNA sequencing with untargeted LC-MS/MS lipidomics provided cohesive mechanistic resolution across both transcriptional and metabolic levels. At the transcriptomic level, DEM reversed the IMQ-induced upregulation of core lipogenic enzymes (*Acly, Cs, Acaca,* and *Hmgc*r) and sustained the repression of key lipid regulators (*Srebf1, Scd1,* and *Acss2*), thereby attenuating pathway activity scores and contracting keratinocyte heterogeneity toward a homeostatic state. Crucially, the persistent quiescence of Acss2, which encodes the primary alternative source of cytosolic acetyl-CoA, indicates that psoriasiform keratinocytes rely heavily on citrate-dependent acetyl-CoA supply and are thus selectively vulnerable to ACLY blockade. At the metabolic level, untargeted lipidomics showed that these transcriptional changes were accompanied by corresponding shifts in the lesional lipidome. DEM counter-regulated 109 IMQ-dysregulated lipid species, driving a coordinated reduction of all 17 canonical de novo synthesized and elongated free fatty acids (C14-C22) and broadly depleting acylcarnitines (84.1% of species reduced), consistent with a reduced supply of fatty acyl chains entering the mitochondrial carnitine shuttle. Functionally, attenuating this pathological lipogenic flux curbed the overproduction of pro-proliferative membrane phospholipids (such as phosphatidylcholines and phosphatidylglycerols) required for rapid keratinocyte division, while simultaneously rescuing IMQ-depleted epidermal barrier lipids, including ceramides, sphingomyelins, and phosphatidylethanolamines. This concordance between single-cell transcriptomics and tissue lipidomics indicates that DEM acts at both the transcriptional and the metabolic level, with the lipid changes in lesional skin running in the direction predicted by ACLY inhibition.

Several limitations should be noted. The polypharmacological nature of DEM means additional targets may contribute to in vivo efficacy. HoloGNN predictions depend on structural model quality, potentially missing poorly characterized targets. The IMQ model, while well-established, does not fully recapitulate chronic relapsing human psoriasis, and translational relevance will require evaluation in complementary models and ultimately clinical settings. In addition, although our data establish ACLY as a direct target and confirm downstream lipidomic reversal in lesional tissue, stable isotope tracing will be valuable in future studies to dynamically quantify carbon flux through the citrate-acetyl-CoA axis, and our findings do not exclude the possibility that parallel targets participate in other context-dependent activities of DEM.

Beyond the specific DEM-ACLY interaction described here, the integrative framework developed in this study may provide a broadly applicable strategy for target identification of bioactive small molecules. Consistent with this, the same screen recovered and biochemically confirmed a second, mechanistically unrelated pair, Chrysosplenol D-ATIC. By combining proteome-wide thermal screening with structure-aware computational prioritization and orthogonal biochemical validation, this workflow offers a practical route to move from candidate stabilization events to direct target assignment. Such an approach should be particularly valuable for structurally complex natural products, for which polypharmacology and uncertain target identity often impede mechanistic interpretation and therapeutic development.

## METHODS

### Training and evaluation of HoloGNN model

The HoloGNN model was trained and evaluated using the PDBbind dataset. Following the data-splitting strategy adopted in GIGN, the model was trained and validated on the PDBbind v2016 dataset, while performance was assessed on three independent benchmark sets: the PDBbind v2013 core set, the PDBbind v2016 core set, and the PDBbind v2019 holdout set. All models were trained three times with different random initializations to ensure robustness. Training was conducted on a single NVIDIA GeForce RTX 3090 GPU with an initial maximum of 800 epochs. The batch size was set to 128 and model parameters were optimized using a learning rate of 5×10^-4^. To mitigate overfitting, a weight decay of 1×10^-6^ was applied, and early stopping was implemented with a patience of 100 epochs, such that training was terminated if the validation loss did not decrease for 100 consecutive epochs. Model performance was assessed using the root mean square error (RMSE) and the Pearson correlation coefficient between predicted and experimental binding affinities. RMSE was used to quantify the prediction error, whereas the Pearson correlation coefficient was used to evaluate the linear correlation between predicted and measured values.

### Molecular docking

Crystal structures of the target proteins were obtained from the PDBbind, including ACLY (PDB ID: 6O0H), BRAF (PDB ID: 6XFP), CSNK2A2 (PDB ID: 6QY9), and CDK11 (PDB ID: 7UKZ). For ACLY, the docking center was defined according to the previously reported active-site region^44^. For the other proteins, docking grids were centered on the positions of the co-crystallized ligands present in the corresponding crystal structures. Docking simulations were performed for all compounds using Uni-Dock^45^, and the resulting docking scores were recorded. For each protein-ligand pair, the binding pose with the highest docking score was selected and used as the structural input for subsequent HoloGNN prediction.

### Selection of experimented natural products

To assemble the set of 50 natural products investigated in this study, we first surveyed multiple curated compound resources, including the NPACT database of plant-derived agents, the TCMSP database supporting traditional Chinese medicine research, along with PubChem, ChemBank, and relevant collections from Selleckchem. These databases were complemented by manual literature curation to identify candidate molecules with reported cellular or in vivo antitumor activity. For each candidate compound, we conducted a systematic search of PubMed and Web of Science for articles published between 2010 and 2024. Based on the collected evidence, including documented bioactivities and associated disease contexts, we assembled a final panel of 50 structurally diverse natural products for screening.

### Cell culture

Human chronic myelogenous leukemia K562 cells (American Type Culture Collection, ATCC) were cultured in RPMI-1640 medium (Gibco) supplemented with 10% fetal bovine serum (FBS; EXCELL). The human immortalized keratinocyte cell line HaCaT was cultured in Dulbecco’s modified Eagle’s medium (DMEM) supplemented with 10% FBS. Both cell lines were maintained at 37 °C in a humidified incubator with a 5% CO_2_ atmosphere. Cells were passaged routinely and harvested for downstream experiments when they reached approximately 80-85% confluence.

### Generation of ACLY-knockdown HaCaT cells

ACLY-knockdown HaCaT cells were generated by lentiviral transduction with shRNAs targeting human ACLY (shACLY-1, 5′-GGGAGGAAGCTGATGAATATA-3′; shACLY-2, 5′-CCCTGAAGACAAGAAAGAAAT-3′), cloned into the pLVX-shRNA2-ZsGreen1-T2A-Puro vector via BamHI/EcoRI sites. A non-targeting scrambled shRNA (shNC, 5′-GTCTCCGAACGTGTCACGT-3′) cloned into the same vector served as control. Transduction efficiency was monitored by ZsGreen1 fluorescence. Transduced cells were selected with puromycin (1 μg/mL) over three passages and maintained as pooled polyclonal populations rather than single-cell clones, in order to avoid clonal bias. Knockdown efficiency was approximately 90% at the protein level as assessed by western blotting, and was verified within the same passage range used for the functional experiments.

### In vitro psoriasis-like inflammatory model

To establish an in vitro psoriasis-like inflammatory model, HaCaT cells were seeded and allowed to adhere overnight, and were then stimulated with the M5 cytokine cocktail (TNF-α, IL-17A, IL-22, IL-1α and Oncostatin-M, 10 ng/mL each; Sino Biological) for 24 h to reproduce the inflammatory and hyperproliferative phenotype of psoriatic keratinocytes. Where cells were subsequently treated with DEM, M5 was maintained in the medium throughout the treatment period and the M5-containing medium was refreshed every 24 h.

### Cell lysis and protein extraction

To prepare cell lysates for downstream biophysical profiling, harvested cells were washed with ice-cold PBS and pelleted. The cell pellets were then resuspended in a non-denaturing lysis buffer consisting of 50 mM HEPES (pH 7.5), 5 mM β-glycerophosphate, 10 mM MgCl_2_, 1 mM tris(2-carboxyethyl)phosphine (TCEP), and a complete protease inhibitor cocktail (Roche). To maximize disruption efficiency while preserving native protein conformations, the cell suspensions were subjected to three sequential cycles of rapid freezing in liquid nitrogen and subsequent thawing in a room-temperature water bath. This was followed by mechanical shearing by ten passages through a 27-gauge syringe needle. The crude lysates were clarified by centrifugation at 21,000 × *g* for 20 min at 4 °C to remove insoluble debris. The total protein concentration of the resulting supernatant was quantified using a bicinchoninic acid (BCA) assay kit (Beyotime). Clarified lysates were aliquoted, snap-frozen, and stored at −80 °C until further analysis.

### Experimental protocol of MAPS-iTSA

A total of 50 natural products (all purchased from MedChemExpress, MCE) were divided into two sets, each distributed into 15 tubes according to the optimized 15×25 sensing matrix. Each compound was pre-dissolved in DMSO at 10 mM. Five drugs were combined in each tube, with each mixture prepared in triplicate, yielding a final concentration of 40 µM per compound in 20 µL containing 2% DMSO. Subsequently, 20 µL of K562 cell lysate (100 µg) was added to each tube and thoroughly mixed, resulting in a final reaction volume of 40 µL containing 20 µM of each compound and 1% DMSO. An additional tube containing methotrexate (MTX) was included as a positive control to ensure experimental quality control. The 16 tubes were then incubated in a PCR thermocycler at 52 °C for 3 min, while two DMSO-only samples were heated at 37 °C for the same duration. After heating, samples were centrifuged at 18,000 rcf for 30 min at 4 °C, and the resulting supernatants were collected for downstream analyses. The complete 50-compound panel was processed twice in this way, on separate days using lysates prepared from two independently thawed and cultured batches of K562 cells; the two runs therefore constitute independent biological replicates of the screen.

### Sample preparation by using SISPROT

A streamlined workflow was implemented for sample preparation^46^. Each 200 µL pipette tip was first packed with a three-layer C18 membrane (3M Empore) and a 1:1 mixture of SCX/SAX beads (final concentration 20 mg/mL). The assembled tips were conditioned sequentially with methanol and potassium citrate buffer (pH 3.0) to ensure full activation. Prior to loading, samples were acidified to pH 2-3 with 1% formic acid and then applied to the prepared tips. The cartridges were rinsed with 8 mM potassium citrate containing 20% acetonitrile, followed by on-tip reduction using 10 mM TCEP for 25 min at room temperature. After a brief wash with 50 mM Tris-HCl (pH 8.0), proteins were digested directly on the tip using trypsin (0.1 µg/µL) and 10 mM iodoacetamide in Tris buffer at 37 °C for 90 min. Peptides were subsequently eluted with 500 mM NaCl. For tandem mass tag (TMT) labeling, the eluate was adjusted to pH 8.0 with 50 mM HEPES, followed by addition of TMT reagent prepared in the same buffer and incubation for 1 h. The labeling reaction was quenched and desalting was performed by adding 1% formic acid. Finally, peptides were extracted with 80% acetonitrile containing 0.5% acetic acid, lyophilized, and stored at -20 °C until LC-MS/MS analysis.

### High-pH reversed-phase fractionation

Peptide separation of the TMT-labeled pooled samples was carried out using a 200 µL pipette tip packed with a twelve-layer C18 disk, which served as the solid-phase carrier for fractionation. The workflow consisted of five major steps. First, the C18 material was conditioned with methanol, rinsed with 1% formic acid, and equilibrated for sample adsorption. Second, the freeze-dried peptides were reconstituted, acidified to pH 2-3 with 1% formic acid, and applied to the C18 tip twice to maximize binding efficiency. Third, the bound peptides were shifted to an alkaline environment using 5 mM ammonium formate to prepare for high-pH fractionation. Fourth, gradient separation was performed by sequentially eluting the peptides with 20 µL of eighteen acetonitrile solutions (3%, 5%, 7%, 9%, 11%, 13%, 15%, 17%, 19%, 21%, 23%, 24%, 26%, 28%, 30%, 35%, 40%, and 80% ACN), all prepared in 5 mM ammonium bicarbonate at pH 10. Finally, the eluates were pooled into six combined fractions, dried in a vacuum concentrator, and preserved at -20 °C for later analysis.

### Extraction of Tripterygium wilfordii

The dried and pulverized *Tripterygium wilfordii* roots were ultrasonically extracted with ethyl acetate at a solid-to-liquid ratio of 1:10 (g/mL) for 30 min. The mixture was filtered, and the extraction process was repeated for three successive cycles. The combined filtrates were concentrated under reduced pressure at 40-45 °C using a rotary evaporator to obtain a crude extract. To completely remove the residual organic solvent, the extract was subsequently dried in a vacuum oven overnight to yield a dried extract. Finally, the extract was weighed to calculate the yield and stored in light-resistant vials at -20 °C until further use.

### LC-MS/MS analysis for MAPS-iTSA

Proteomic measurements were carried out using a Thermo Fisher Scientific U3000 HPLC system interfaced with an Orbitrap Exploris 480 mass spectrometer. Chromatographic separation employed a binary solvent system, with solvent A being water containing 0.1% formic acid and solvent B consisting of 100% acetonitrile supplemented with 0.1% formic acid. Peptide samples were reconstituted in 0.1% formic acid and loaded onto an integrated emitter column (100 μm inner diameter, 20 cm length) packed with 1.9 μm, 120 Å ReproSil-Pur C18 material (Dr. Maisch GmbH). Peptide elution was performed over a 120-min high-resolution gradient, starting with an increase from 6% to 10% B in the first 2 minutes, followed by a gradual shift to 28% B over 95 minutes, then rising to 40% B within 11 minutes, and finally ramping to 99% B in the last 2 minutes. A 3-minute wash at 99% B and a subsequent 7-minute re-equilibration at 1% B completed the run. The flow rate was kept constant at 500 nL/min. The Orbitrap Exploris 480 operated under data-dependent acquisition (DDA), collecting one full MS survey scan prior to selecting the 50 most intense precursors for fragmentation in each cycle. When turbo-TMT was enabled, MS1 spectra were recorded at 60,000 resolution and MS/MS spectra at 30,000 resolution. The acquisition window spanned 350-1,200 m/z, with a maximum ion injection time of 45 ms. Dynamic exclusion was set for 45 s, and precursor selection used a 0.7 m/z isolation width with an HCD collision energy of 38%.

TMT-acquired mass spectrometry data were analyzed using Proteome Discoverer software (v2.4, Thermo Fisher Scientific). Data processing employed the curated human UniProt FASTA database (downloaded May 23, 2024) with trypsin specified as the enzyme and up to 2 missed cleavages allowed, a precursor ion mass tolerance of 20 ppm and a fragment ion mass tolerance of 0.02 Da. Carbamidomethylation of cysteine and TMT labelling of peptide N-termini and lysine residues were set as fixed modifications, and oxidation of methionine and protein N-terminal acetylation as variable modifications. The false discovery rate was restricted to 1% at both the peptide-spectrum match and peptide identification levels. Reporter ion measurements were filtered using a 50% co-isolation cutoff to ensure accurate quantification.

### Western blotting

Equal amounts of protein samples were subjected to electrophoresis on 12.5% SDS-polyacrylamide gels and subsequently transferred onto polyvinylidene difluoride (PVDF) membranes (Millipore, USA). After transfer, membranes were blocked with 1× protein-free rapid blocking buffer (Epizyme, China) for 15 min at room temperature. The membranes were then incubated overnight at 4 °C with gentle agitation using primary antibodies against ACLY (Proteintech, 67166-1-Ig; 1:1000) and GAPDH (Proteintech, 60004-1-Ig; 1:50000). Following primary antibody incubation, the membranes were washed and probed with horseradish peroxidase (HRP)-conjugated secondary antibodies (Proteintech, China; 1:5000) for 1 h at room temperature. Protein signals were detected using an enhanced chemiluminescence (ECL) kit (Bio-Rad, USA) according to the manufacturer’s instructions.

### Histone acetylation as a read-out of cellular ACLY function

HaCaT cells were seeded in six-well plates (2 × 10 cells per well), grown overnight to 70-80% confluence and treated for 24 h with vehicle (0.1% DMSO), DEM (5, 10 or 20 μM), DEM (20 μM) plus 5 mM sodium acetate, 5 mM sodium acetate alone, or 5 mM NaCl; DMSO was 0.1% in all conditions.

Cells were washed twice on ice with cold PBS containing deacetylase inhibitors (10 mM sodium butyrate, 5 mM nicotinamide, 1 μM trichostatin A) and lysed directly in 150 μL of 1× Laemmli buffer containing the same inhibitors and a protease inhibitor cocktail; the whole lysate was carried forward without clearing centrifugation. Lysates were sonicated on ice and heated at 95 °C for 10 min. Equal volumes were resolved on 15% SDS-PAGE gels, transferred to 0.2-μm PVDF membranes and blocked with 1× protein-free rapid blocking buffer (Epizyme, China). Membranes were probed overnight at 4 °C in 5% BSA/TBST with antibodies against H3K9ac (CST, #9649; 1:1000), H3K27ac (CST, #8173; 1:1000) and total histone H3 (CST, #4499; 1:2000), followed by an HRP-conjugated secondary antibody (1:5000) and ECL detection at non-saturating exposures. Band intensities were quantified in ImageJ, normalized to total histone H3 in the same lane and expressed relative to the vehicle control of the same experiment. Data are from three independent biological replicates performed on separate days, with all conditions of a replicate resolved on the same gel. Significance was assessed by one-way ANOVA with Dunnett’s post hoc test against vehicle, and the acetate rescue by a pre-specified comparison between DEM alone and DEM plus acetate.

### Cellular thermal shift assay

CETSA was performed in the cell-lysate format. HaCaT cell lysates were incubated with DEM (50 μM) or an equivalent volume of DMSO at 37 °C for 10 min. Equal aliquots of lysate were distributed into PCR tubes and heated at 37, 43, 46, 49, 52, and 55 °C for 3 min in a PCR thermocycler. After cooling to room temperature, samples were centrifuged at 20,000 × g for 20 min at 4 °C to remove denatured and precipitated proteins. Soluble fractions were analyzed by western blotting with anti-ACLY and anti-GAPDH antibodies as described above. Band intensities were quantified using ImageJ and normalized to GAPDH. The same lysate-format procedure was applied to the Chrysosplenol D_-_ATIC pair using AC16 cell lysates (Procell) incubated with Chrysosplenol D (50 μM) or DMSO, with heating at 37, 42, 46, 49, 52, 55 and 58 °C; soluble endogenous ATIC was detected with anti-ATIC (Proteintech, cat. no. 10726-1-AP; 1:2000) and normalized to β-actin (Proteintech, cat. no. 66009-1-Ig; 1:20000) in the same lane and to the 37 °C DMSO control. All experiments were performed in three independent biological replicates (n = 3).

### Surface plasmon resonance assay

SPR was performed on a Biacore 8K system (Cytiva). The purified recombinant human ACLY used for SPR and DARTS was the same full-length, GST-tagged protein as in the enzymatic assay (BPS Bioscience, cat. no. 50255; Sf9 insect cells). ACLY was buffer-exchanged into 10 mM sodium acetate (pH 4.0) on a desalting column, diluted to 40 μg/mL and immobilized to ∼11,000 RU by standard amine coupling. Binding was measured at 25 °C in PBST containing 1% DMSO at 30 μL/min, with a 90 s association and 180 s dissociation phase; DEM was injected at eight concentrations in a two-fold series (1.56-200 μM). Data were analysed in Biacore Insight Evaluation Software using a steady-state affinity model.The Chrysosplenol D-ATIC interaction was measured under the same conditions using purified recombinant human ATIC (full-length, residues 1-592, N-terminally GST-tagged, Escherichia coli; ipodix, cat. no. PA2000-5677), reconstituted in ddH2O, buffer-exchanged as above. Chrysosplenol D was injected at six concentrations (200, 100, 50, 25, 12.5 and 5 μM).

### Drug affinity responsive target stability assay

The same GST-tagged recombinant human ACLY described above (2 μg per reaction) was incubated with DEM (20 μM) or DMSO (final DMSO ≤ 0.1%) in reaction buffer (50 mM HEPES pH 7.5, 150 mM NaCl, 10 mM MgCl) for 1 h at room temperature, then digested with pronase (Roche) at enzyme-to-protein mass ratios of 1:2000, 1:1000, 1:500 and 1:100 for 20 min at 25 °C, with an undigested control (no pronase) for each condition; reactions were stopped by boiling in SDS loading buffer. Band intensities were quantified by densitometry in ImageJ and normalized to the corresponding undigested control (n = 3). For the Chrysosplenol D-ATIC pair, DARTS was performed in the same way on AC16 cell lysates (Procell), using Chrysosplenol D (50 μM) and pronase ratios of 1:1000, 1:500 and 1:100; residual endogenous ATIC was detected with the anti-ATIC antibody described above (n = 4).Significance between compound- and DMSO-treated samples at each pronase ratio was assessed by two-tailed unpaired Student’s t-test.

### ACLY enzymatic activity assay

ACLY enzymatic activity was measured with the ACLY Assay Kit (BPS Bioscience, cat. no. 79904), which supplies the purified recombinant human ACLY used in this assay (cat. no. 50255), substrate, ATP and assay buffer, with ADP-Glo Kinase Assay reagent (Promega, cat. no. V6930) for detection. All conditions were performed in three independent experiments (n = 3). Assays were assembled in 25-μL reactions containing ATP, Coenzyme A, sodium citrate, and ACLY assay buffer. Test inhibitors were prepared as 100× DMSO stocks, diluted in water, and added to the designated wells; vehicle controls received an equivalent volume of inhibitor buffer. Final DMSO content in all reactions was maintained at ≤1%. Blank wells contained assay buffer in place of enzyme. Reactions were initiated by adding diluted ACLY enzyme (4 ng/μL) and incubated at 30 °C for 45 min. ADP-Glo reagent was then added to terminate ATP consumption and deplete unreacted ATP, followed by a 45-min incubation at room temperature. Subsequently, Kinase Detection reagent was added to convert ADP to ATP and generate a luminescent signal proportional to ACLY activity after a further 45-min incubation. Luminescence was recorded using a microplate luminometer in chemiluminescence mode with a 1-s integration time and top-read optics. Blank readings were subtracted from all values prior to analysis.

### In vitro cell proliferation assay

For dose-response analysis in ACLY-knockdown cells, shNC, shACLY-1 and shACLY-2 HaCaT cells were seeded in 96-well plates, stimulated with the M5 cytokine cocktail for 24 h as described above, and then treated with serially diluted DEM or vehicle (0.1% DMSO) in fresh M5-containing medium for a further 72 h, with the medium refreshed every 24 h. Inhibition was calculated relative to the vehicle-treated wells of the same cell line, so that the lower basal proliferation rate of ACLY-knockdown cells does not confound the comparison. IC values were obtained by fitting a four-parameter log(inhibitor)-response model and are reported as the mean of three independent biological experiments.

### In vivo psoriasis mouse model

Female C57BL/6 mice (6-8 weeks) were randomly allocated into strictly defined groups using a simple randomization method (n = 5 mice per group): Control, IMQ, and IMQ+DEM. No statistical methods were used to predetermine sample sizes, but our sample sizes are consistent with standards generally employed in similar in vivo studies. Psoriasis-like dermatitis was induced by daily topical application of 62.5 mg of 5% IMQ cream on the shaved dorsal skin for 4 consecutive days. The IMQ+DEM group concurrently received daily intraperitoneal injections of DEM (1 mg/kg). Skin inflammation was blindly scored daily by two independent researchers who were completely unaware of the group allocation, according to the clinical Psoriasis Area and Severity Index (PASI) criteria (erythema, scaling, and thickness; scale 0-4). At the end of the treatment period, mice were euthanized for tissue harvesting. Tissues were fixed in 4% paraformaldehyde, sectioned (4-5 μm), and H&E stained to evaluate skin pathology (hyperkeratosis, acanthosis) and internal organ toxicology. Pathological scoring differences between groups were evaluated by two-tailed unpaired Student’s t-test. No animals or data points were excluded from the analysis.

### qPCR analysis

Total RNA was extracted from skin tissues using TRIzol reagent. RNA was reverse-transcribed into cDNA using a reverse transcription kit. Quantitative real-time PCR was performed with SYBR Green master mix, using β-actin as the housekeeping gene. The relative mRNA expression levels of pro-inflammatory cytokines (Ccl3, Il6, and Tnf) were determined by the Livak method. Relative expression was compared between groups by two-tailed unpaired Student’s t-test.

### Single-cell suspension preparation

Dorsal skin from NC, IMQ, and IMQ+DEM mice was minced (∼0.5 mm³) and enzymatically digested (MACS Tumor Dissociation Kit, Miltenyi Biotec; 37°C, 30 min). Cell suspensions were filtered through 70-μm and 40-μm strainers (BD Biosciences) and centrifuged (300 × g, 10 min, 4°C). Erythrocytes were lysed (RBC Lysis Buffer, Thermo Fisher Scientific), and pelleted cells were washed and resuspended in PBS with 0.04% BSA. Viability exceeded 85% by Trypan Blue exclusion prior to library construction.

### scRNA-seq library construction and sequencing

scRNA-seq libraries were constructed by BGI Genomics using the DNBelab C Series High-throughput Single-Cell RNA Library Preparation Set (MGI, #940-000519-00). Encapsulated single cells underwent emulsion breakage, mRNA capture, and reverse transcription. cDNA was amplified, fragmented (300-500 bp), and indexed. Libraries were quantified via Qubit ssDNA Assay (Thermo Fisher Scientific) and Agilent 2100 Bioanalyzer, and sequenced on the DNBSEQ-T7 platform (BGI-Shenzhen) with paired-end reads (Read 1: 30 bp for dual 10-bp cell barcodes and 10-bp UMI (unique molecular identifier); Read 2: 100 bp gene insert).

### Affinity ultrafiltration mass spectrometry assay

The incubation mixture was prepared by combining 100 μL of a 400 μg/mL *Tripterygium wilfordii* solution with 100 μL of ACLY (100 μg/mL) in PBS (pH 7.8) to achieve a final volume of 200 μL. After incubation at 37 °C for 30 min, the mixture was transferred to an ultrafiltration unit equipped with a membrane designed to retain molecules above 10 kDa MW, followed by centrifugation at 12,000g for 20 min at 4 °C. The membrane was washed four times with 200 μL of PBS (pH 7.8) to remove unattached molecules. Subsequently, complexes of ACLY and its ligands were moved a new tube, and ligands were dissociated from ACLY by ultrasonication in 200 μL of methanol for 20 min. Ligands freed from ACLY were isolated by centrifugation at 12,000 g for 15 min at 4 °C, after which the supernatant was collected and evaporated to dryness under vacuum.The residues were reconstituted in 100 μL of methanol and subjected to final centrifugation at 12,000g for 15 min at 4 °C prior to UHPLC-MS/MS analysis. Data analysis was assisted by XCMS^47^ and DeepMASS^48^. Then, identified metabolites were confirmed with standards.

### Untargeted LC-MS/MS lipidomic profiling

Dorsal skin tissues from NC, IMQ, and IMQ+DEM mice (n = 3 per group) were collected and snap-frozen in liquid nitrogen. Untargeted lipidomic profiling was performed by Novogene Co., Ltd. (Beijing, China) using liquid chromatography-tandem mass spectrometry (LC-MS/MS). Lipids were extracted using a standard methyl tert-butyl ether (MTBE)/methanol/water biphasic protocol and reconstituted for LC-MS/MS analysis in both positive and negative electrospray ionization modes. Pooled quality control (QC) samples were injected periodically throughout the run to assess instrumental stability. Raw data files (.raw) were processed using LipidSearch for peak extraction, retention time alignment, and lipid identification based on precursor m/z, mass tolerance, adduct forms, and MS/MS spectral matching. Peak areas were integrated via XCMS after format conversion (.mzXML) using msconvert, and normalized to total peak area. Lipid features with a coefficient of variation (CV) > 30% across QC samples were removed, leaving 1,271 positive-mode and 640 negative-mode species (1,911 lipids across 26 subclasses) for downstream analysis.

### Lipidomic statistical and pathway analysis

Multivariate analyses, including principal component analysis (PCA) and partial least squares discriminant analysis (PLS-DA), were conducted using metaX software. Differential lipids were screened using fold change (FC > 1.2 or < 0.833), variable importance in the projection (VIP > 1.0), and two-tailed Student’s t-test (P < 0.05). Functional annotations were mapped against KEGG, HMDB, and LIPID MAPS databases.

### Data processing, clustering, and metabolic scoring

Raw data were processed via DNBelab_C_Series_scRNA-analysis-software and aligned to the mm10 genome using STAR (v2.5.1b)^49^ to generate expression matrices. Downstream analysis utilized Seurat (v5.1.0)^50^. Cells were stringently filtered (500-5,000 genes, ≤20,000 UMIs, ≤15% mitochondrial reads), yielding 44,744 high-quality cells (NC: 10,639; IMQ: 24,224; IMQ+DEM: 9,881). Data were log-normalized, and the top 2,000 highly variable genes were used for Principal Component Analysis (PCA). Cell populations were clustered (FindClusters) and visualized via UMAP. Keratinocyte-specific biological processes were identified by GSEA using the cluster Profiler^51^. ACLY-dependent lipid metabolic reprogramming was evaluated using Seurat’s AddModuleScore function with a targeted gene set (*Acly, Fasn, Scd1, Srebf1, Srebf2, Hmgcr, Acss2, Slc25a1*). ACLY pathway scores were compared between the IMQ and IMQ+DEM groups using Welch’s t-test. The spatial distribution of the pathway scores among cell populations was visualized using UMAP feature plots. Additionally, macroscopic inter-group score variations were depicted via violin plots, while the detailed expression dynamics of individual ACLY pathway genes across the experimental groups were illustrated using dot plots.

## ETHICS STATEMENT

This study did not involve human participants. All animal experiments were approved by the Ethics Committee of Xiangya Hospital, Central South University, China (approval number: 202310046), and all procedures were conducted in accordance with the ethical guidelines for animal research of Xiangya Hospital, Central South University, China.

## DATA AVAILABILITY

The raw mass spectrometry proteomics data have been deposited in the ProteomeXchange Consortium via the PRIDE partner repository under the dataset identifier PXD086106. Processed proteomics data are provided in the Supplementary Information. All other data supporting the findings of this study are released together with the code.

## CODE AVAILABILITY

The source code, readme file, along with minimum example and datasets are available at https://github.com/hcji/NPTD.

## ACKNOWLEDGMENTS

This work is financially supported by Guangdong Basic and Applied Basic Research Foundation (Grant No. 2025A1515012831), and the National Natural Science Foundation of China (Grant No. 32470685, 82606899).

## REFERENCES

1. Harvey, A. L., Edrada-Ebel, R. & Quinn, R. J. The re-emergence of natural products for drug discovery in the genomics era. Nat Rev Drug Discov 14, 111–129 (2015).

2. Atanasov, A. G., Zotchev, S. B., Dirsch, V. M. & Supuran, C. T. Natural products in drug discovery: advances and opportunities. Nat. Rev. Drug Discovery 20, 200–216 (2021).

3. Jin, C. et al. Synthesis and biological evaluation of paclitaxel and camptothecin prodrugs on the basis of 2-nitroimidazole. ACS Med. Chem. Lett. 8, 762–765 (2017).

4. Dai, L. et al. Target identification and validation of natural products with label-free methodology: A critical review from 2005 to 2020. Pharmacology & Therapeutics 216, 107690 (2020).

5. Lomenick, B. et al. Target identification using drug affinity responsive target stability (DARTS). Proc Natl Acad Sci U S A 106, 21984–21989 (2009).

6. Feng, Y. et al. Global analysis of protein structural changes in complex proteomes. Nat Biotechnol 32, 1036–1044 (2014).

7. Strickland, E. C. et al. Thermodynamic analysis of protein-ligand binding interactions in complex biological mixtures using the stability of proteins from rates of oxidation. Nat Protoc 8, 148–161 (2013).

8. Savitski, M. M. et al. Tracking cancer drugs in living cells by thermal profiling of the proteome. Science 346, 1255784 (2014).

9. Dziekan, J. M. et al. Cellular thermal shift assay for the identification of drug–target interactions in the Plasmodium falciparum proteome. Nat Protoc 15, 1881–1921 (2020).

10. Ball, K. A. et al. An isothermal shift assay for proteome scale drug-target identification. Commun Biol 3, 75 (2020).

11. Ji, H. et al. Target deconvolution with matrix-augmented pooling strategy reveals cell-specific drug-protein interactions. Cell Chemical Biology 30, 1478–1487.e7 (2023).

12. Tan, C. S. H. et al. Thermal proximity coaggregation for system-wide profiling of protein complex dynamics in cells. Science 359, 1170–1177 (2018).

13. Kozoriz, K. & Lee, J.-S. Chemical proteomics for a comprehensive understanding of functional activity and the interactome. Chemical Society Reviews 54, 6186–6207 (2025).

14. Sim, J., Kim, D., Kim, B., Choi, J. & Lee, J. Recent advances in AI-driven protein-ligand interaction predictions. Current Opinion in Structural Biology 92, 103020 (2025).

15. Öztürk, H., Özgür, A. & Ozkirimli, E. DeepDTA: deep drug–target binding affinity prediction. Bioinformatics 34, i821–i829 (2018).

16. Li, S. et al. MONN: A Multi-objective Neural Network for Predicting Compound-Protein Interactions and Affinities. Cell Systems 10, 308–322.e11 (2020).

17. Wan, F., et al. DeepCPI: A Deep Learning-Based Framework for Large-Scale *in Silico* Drug Screening. Genomics, Proteomics & Bioinformatics 17, 478–495 (2019).

18. CarsiDock: a deep learning paradigm for accurate protein–ligand docking and screening based on large-scale pre-training. Chem. Sci. 15, 1449–1471 (2024).

19. Zhang, X. et al. Efficient and accurate large library ligand docking with KarmaDock. Nat. Comput. Sci. 3, 789–804 (2023).

20. Nehil-Puleo, K., Quach, C. D., Craven, N. C., McCabe, C. & Cummings, P. T. E(n) equivariant graph neural network for learning interactional properties of molecules. J. Phys. Chem. B 128, 1108–1117 (2024).

21. Yang, Z., Zhong, W., Lv, Q., Dong, T. & Yu-Chian Chen, C. Geometric interaction graph neural network for predicting protein–ligand binding affinities from 3D structures (GIGN). J. Phys. Chem. Lett. 14, 2020–2033 (2023).

22. Sun, X. et al. Therapeutic potential of demethylzeylasteral, a triterpenoid of the genus Tripterygium wilfordii. Fitoterapia 163, 105333 (2022).

23. Li, Y. et al. Demethylzeylasteral inhibits proliferation, migration, and invasion through FBXW7/c Myc axis in gastric cancer. MedComm (2020) 2, 467–480 (2021).

24. Batchuluun, B., Pinkosky, S. L. & Steinberg, G. R. Lipogenesis inhibitors: Therapeutic opportunities and challenges. Nat Rev Drug Discov 21, 283–305 (2022).

25. Gautam, J. et al. ACLY inhibition promotes tumour immunity and suppresses liver cancer. Nature 645, 507–517 (2025).

26. Kao, Y.-S. et al. Metabolic reprogramming of interleukin-17-producing γδ T cells promotes ACC1-mediated de novo lipogenesis under psoriatic conditions. Nat Metab 7, 966–984 (2025).

27. Lin, A. et al. Off-target toxicity is a common mechanism of action of cancer drugs undergoing clinical trials. Sci. Transl. Med. 11, eaaw8412 (2019).

28. Ru, Y. et al. Role of keratinocytes and immune cells in the anti-inflammatory effects of *Tripterygium wilfordii* Hook. f. in a murine model of psoriasis. Phytomedicine 77, 153299 (2020).

29. Yu, H. et al. Integrated Thermal Proteome Profiling and Affinity Ultrafiltration Mass Spectrometry (iTPAUMS): A Novel Paradigm for Elucidating the Mechanism of Action of Natural Products. Anal. Chem. 96, 15980–15990 (2024).

30. Xiang, W. et al. Inhibition of ACLY overcomes cancer immunotherapy resistance via polyunsaturated fatty acids peroxidation and cGAS-STING activation. Sci. Adv. 9, eadi2465 (2023).

31. Liang, J.-J. et al. Recent advance of ATP citrate lyase inhibitors for the treatment of cancer and related diseases. Bioorganic Chemistry 142, 106933 (2024).

32. Granchi, C. ATP-citrate lyase (ACLY) inhibitors as therapeutic agents: a patenting perspective. Expert Opinion on Therapeutic Patents 32, 731–742 (2022).

33. Nguyen, P. T. T., et al. Acetyl-CoA synthesis in the skin is a key determinant of systemic lipid homeostasis. Cell Reports 44, (2025).

34. Wei, X., Schultz, K., Bazilevsky, G. A., Vogt, A. & Marmorstein, R. Molecular basis for acetyl-CoA production by ATP-citrate lyase. Nat Struct Mol Biol 27, 33–41 (2020).

35. Li, R. et al. ATP-citrate lyase controls endothelial gluco-lipogenic metabolism and vascular inflammation in sepsis-associated organ injury. Cell Death Dis 14, 401 (2023).

36. Lu, Y. et al. ACLY-induced reprogramming of glycolytic metabolism plays an important role in the progression of breast cancer. ABBS 55, 878–881 (2023).

37. Saudou, F. ACLY links mutant α-synuclein to metabolism, autophagy and neurodegeneration. Neuron 113, 1847–1849 (2025).

38. Sun, Q. et al. Bempedoic Acid Unveils Therapeutic Potential in Non-Alcoholic Fatty Liver Disease: Suppression of the Hepatic PXR-SLC13A5/ACLY Signaling Axis. Drug Metab Dispos 51, 1628–1641 (2023).

39. Duarte Lau, F. & Giugliano, R. P. Adenosine Triphosphate Citrate Lyase and Fatty Acid Synthesis Inhibition: A Narrative Review. JAMA Cardiol 8, 879–887 (2023).

40. Zhang, Y. et al. Demethylzeylasteral induces PD-L1 ubiquitin-proteasome degradation and promotes antitumor immunity via targeting USP22. Acta Pharm Sin B 14, 4312–4328 (2024).

41. Pan, L. et al. Demethylzeylasteral targets lactate by inhibiting histone lactylation to suppress the tumorigenicity of liver cancer stem cells. Pharmacol Res 181, 106270 (2022).

42. Conte, A. et al. Advanced morphological control over Cu nanowires through a design of experiments approach. Mater Adv 5, 8836–8846 (2024).

43. Granchi, C. ATP-citrate lyase (ACLY) inhibitors as therapeutic agents: a patenting perspective. Expert Opin Ther Pat 32, 731–742 (2022).

44. Liang, J.-J. et al. Recent advance of ATP citrate lyase inhibitors for the treatment of cancer and related diseases. Bioorganic Chemistry 142, 106933 (2024).

45. Yu, Y. et al. Uni-Dock: GPU-Accelerated Docking Enables Ultralarge Virtual Screening. J. Chem. Theory Comput. 19, 3336–3345 (2023).

46. Chen, W. et al. Simple and Integrated Spintip-Based Technology Applied for Deep Proteome Profiling. Anal Chem 88, 4864–4871 (2016).

47. Forsberg, E. M. et al. Data processing, multi-omic pathway mapping, and metabolite activity analysis using XCMS Online. Nat Protoc 13, 633–651 (2018).

48. Jiang, S. et al. DeepMASS v.2: An enhanced deep learning platform for large-scale discovery and structural annotation of unknown plant metabolites. Plant Communications 101976 (2026) doi:10.1016/j.xplc.2026.101976.

49. Dobin, A. et al. STAR: ultrafast universal RNA-seq aligner. Bioinformatics 29, 15–21 (2013).

50. Hao, Y. et al. Dictionary learning for integrative, multimodal and scalable single-cell analysis. Nat Biotechnol 42, 293–304 (2024).

51. Wu, T. et al. clusterProfiler 4.0: A universal enrichment tool for interpreting omics data. Innovation (Camb) 2, 100141 (2021).

