## Supplementary material for "Thermal Proteomics and AI-assisted Target Deconvolution Identify ACLY as a Direct Target of Demethylzeylasteral in Psoriasis": Supp. Figures S1-S4

**Table of content**

Fig.S1. Benchmarking of HoloGNN against established protein-ligand binding affinity prediction models.

Fig.S2. Recovery and biochemical validation of a second compound-target pair, Chrysosplenol D-ATIC.

Fig.S3. Single-cell transcriptomic characterization of skin cell lineages and keratinocyte subpopulations.

Fig.S4. Untargeted LC-MS/MS lipidomics reveals global counter-regulation of the psoriatic lipidome by Demethylzeylasteral.

**Tables S1-S7 are provided as a separate Excel file.**

Table S1. List of drugs with the sensing matrix of the MAPS-iTSA experiment.

Table S2. Quantitative protein table of MAPS-iTSA experiment I group I.

Table S3. Quantitative protein table of MAPS-iTSA experiment I group II.

Table S4. Quantitative protein table of MAPS-iTSA experiment II group I.

Table S5. Docking scores and HoloGNN-predicted binding affinities of the 50 natural products against ACLY.

Table S6. Quantitative protein table of iTSA of *Tripterygium wilfordii* extract in HepG2 cells.

Table S7. Quantitative protein table of iTSA of *Tripterygium wilfordii* extract in THP-1 cells.

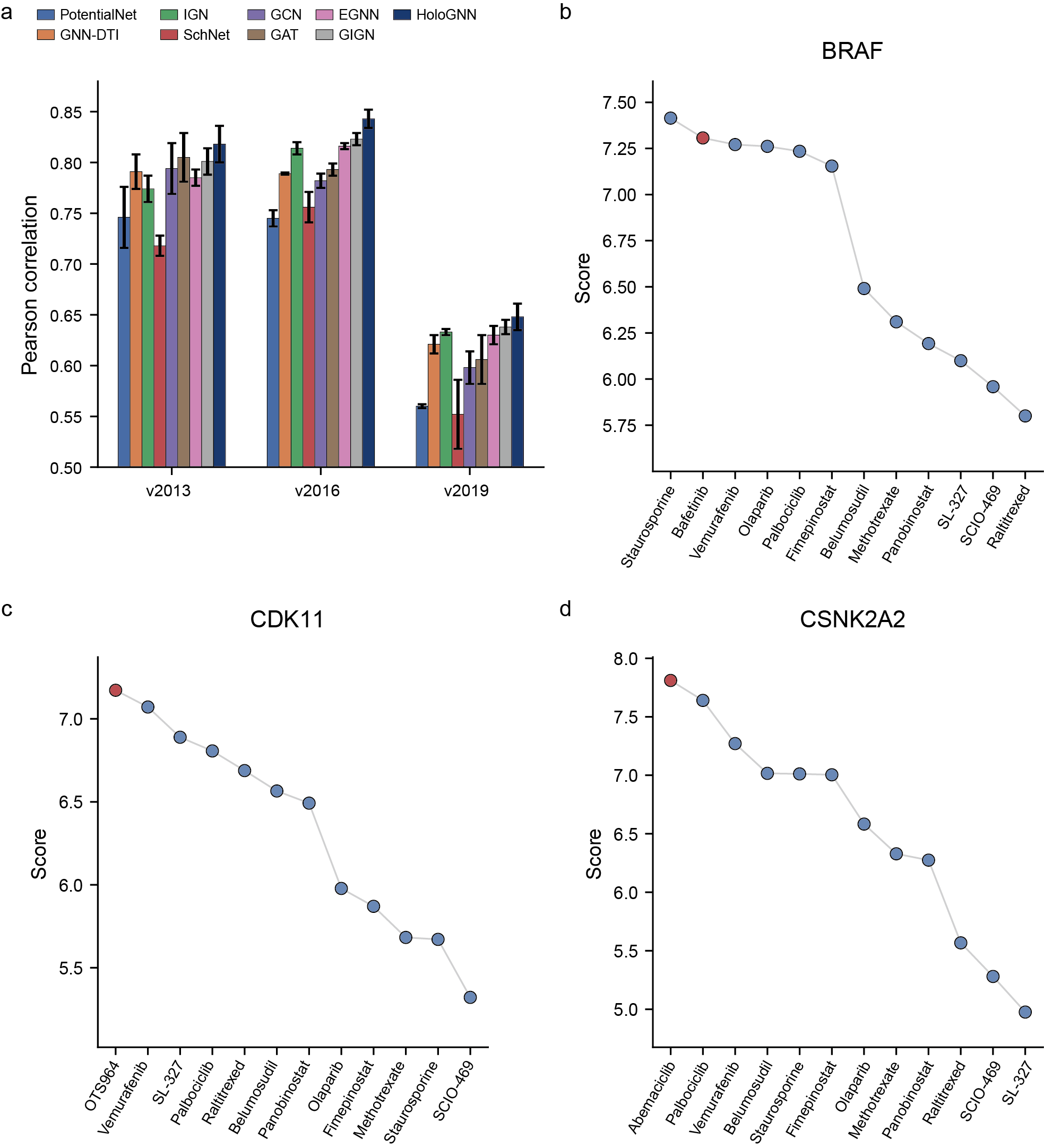

**Fig.S1. Benchmarking of HoloGNN against established protein-ligand binding affinity prediction models. a.** Pearson correlation between predicted and experimental binding affinities for HoloGNN and eight published models on three PDBbind benchmark sets (v2013 and v2016 core sets; v2019 holdout set). Bars are mean ± s.d. of three training runs with different random initializations. **b-d.** Compounds ranked by HoloGNN-predicted binding score against BRAF (b), CDK11 (c) and CSNK2A2 (d); the reported ligand of each target is shown in red.
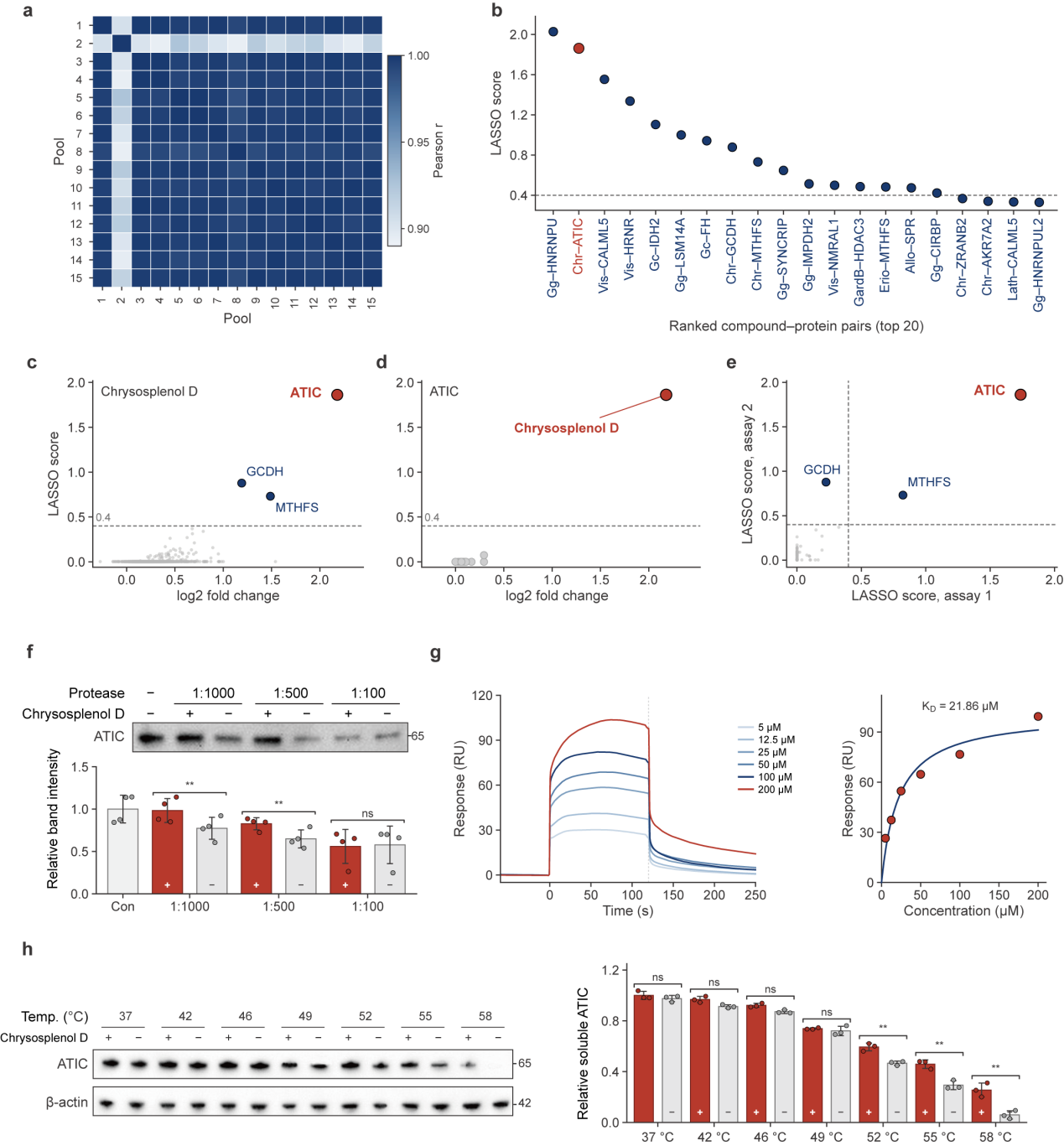

**Fig.S2. Recovery and biochemical validation of a second compound-target pair, Chrysosplenol D-ATIC. a.** Pairwise Pearson correlation of protein abundance between the 15 compound pools of the MAPS-iTSA screen. **b.** Top 20 compound-protein pairs ranked by LASSO score; dashed line, 0.4 selection threshold. **c, d.** LASSO score against log₂ fold change for the candidate proteins of Chrysosplenol D (c) and the candidate compounds of ATIC (d). **e.** LASSO scores of the candidate proteins for Chrysosplenol D in the two independent screens; dashed lines, 0.4 threshold on each axis. **f.** DARTS assay of ATIC protection against pronase digestion. Top, representative blots at the indicated protease-to-protein mass ratios (+/−, presence or absence of Chrysosplenol D); bottom, band intensity normalized to the no-protease control (Con). **g.** SPR analysis of Chrysosplenol D binding to immobilized ATIC. Left, sensorgrams at 5-200 μM; right, steady-state affinity fit (K_D_ = 21.86 μM). **h.** CETSA (cell-lysate format) of endogenous ATIC incubated with Chrysosplenol D (+) or DMSO (−). Left, representative blots; right, soluble ATIC normalized to β-actin in the same lane and to the 37 °C DMSO control. Data are mean ± s.e.m. of four (f) or three (h) independent biological replicates; two-tailed unpaired Student's t-test. ns, not significant; **P < 0.01. DARTS, drug affinity responsive target stability; SPR, surface plasmon resonance; LASSO, least absolute shrinkage and selection operator; DMSO, dimethyl sulfoxide. Compound abbreviations in b: Gg, (−)-Gallocatechin gallate; Chr, Chrysosplenol D; Vis, Visnagin; Gc, (−)-Gallocatechin; GardB, Gardenin B; Erio, Eriocitrin; Allo, Alloimperatorin; Lath, Lathyrol.

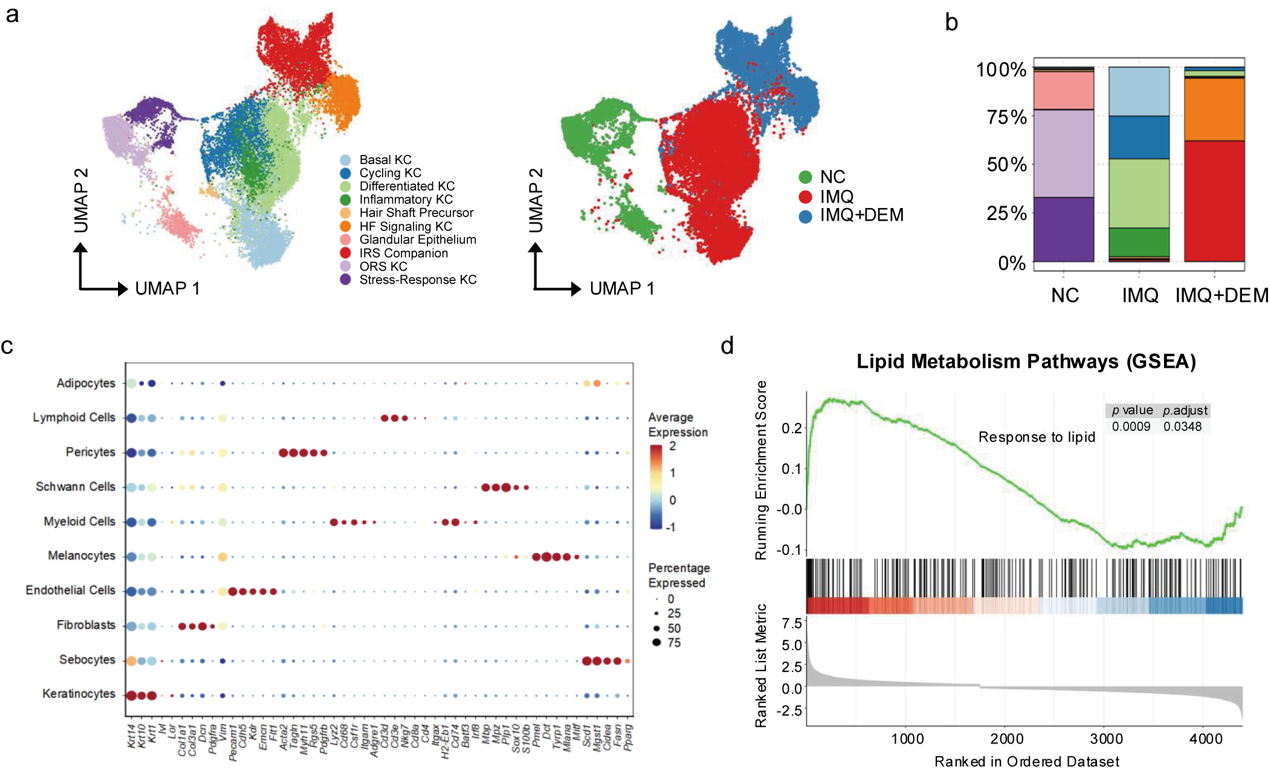

**Fig.S3. Single-cell transcriptomic characterization of skin cell lineages and keratinocyte subpopulations. a.** UMAP visualization of keratinocyte (KC) subtypes, colored by subtype (left) and by group (right). **b.** Relative proportion of KC subtypes across the NC, IMQ and IMQ+DEM groups. **c.** Dot plot showing the expression of canonical lineage marker genes across all identified skin cell types. Dot size represents the percentage of cells expressing the gene, and color intensity indicates average expression level. **d.** Gene set enrichment analysis (GSEA) plot showing significant positive enrichment of lipid metabolism pathways ("Response to lipid", P = 0.0009, adjusted P = 0.0348) in IMQ-stimulated keratinocytes. NC, normal control; IMQ, imiquimod; DEM, Demethylzeylasteral; UMAP, uniform manifold approximation and projection; HF, hair follicle; IRS, inner root sheath; ORS, outer root sheath.

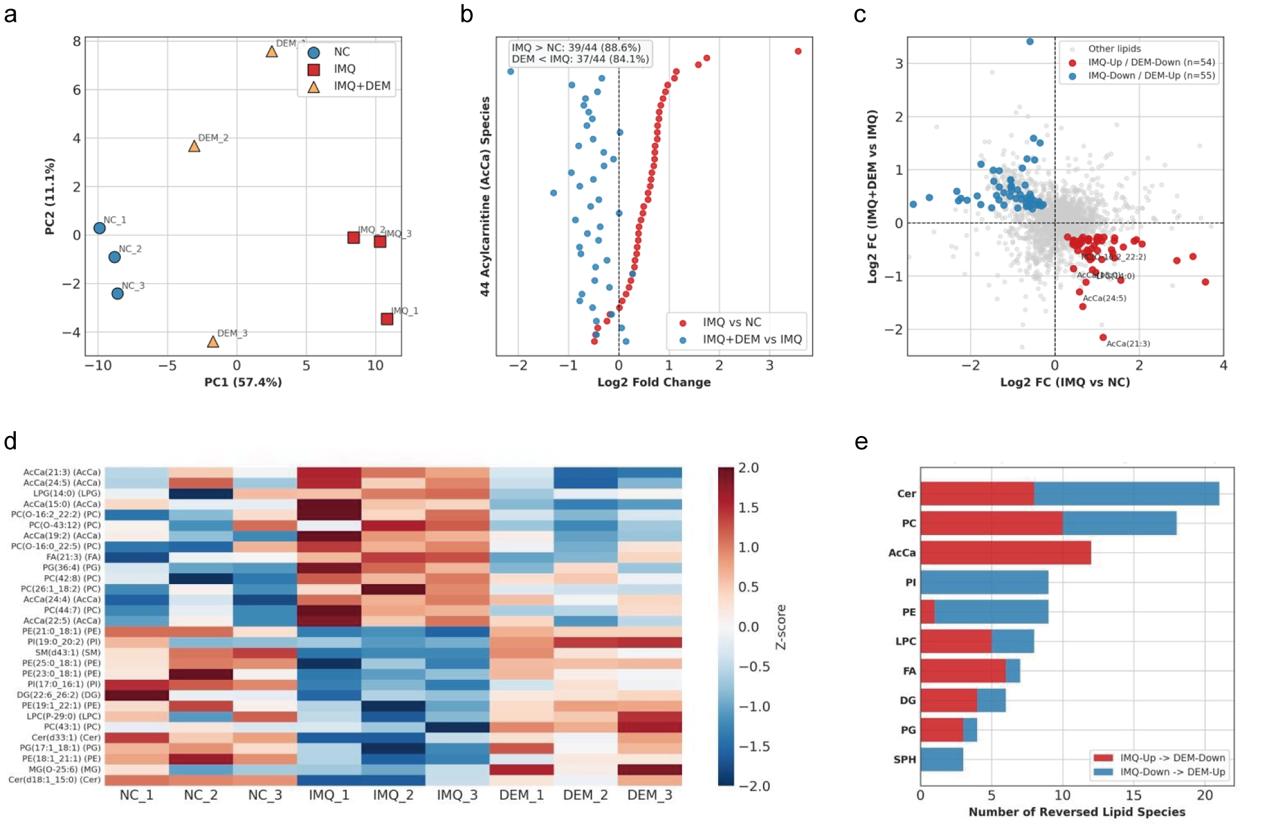

**Fig.S4. Untargeted LC-MS/MS lipidomics reveals global counter-regulation of the psoriatic lipidome by Demethylzeylasteral. a.** Principal component analysis (PCA) score plot of the Demethylzeylasteral-responsive lipidome (109 lipid species) across NC, IMQ, and IMQ+DEM groups (n = 3 biologically independent mice per group). **b.** Bidirectional fold-change distribution of all 44 detected acylcarnitine (AcCa) species, showing global accumulation in IMQ versus NC (red, 39/44 species upregulated, 88.6%) and widespread suppression in IMQ+DEM versus IMQ (blue, 37/44 species downregulated, 84.1%). **c.** Four-quadrant scatter plot of log_2_ fold changes comparing IMQ versus NC (x-axis) against IMQ+DEM versus IMQ (y-axis) across all 1,911 quantified lipids, highlighting 54 IMQ-Up/DEM-Down species (red) and 55 IMQ-Down/DEM-Up species (blue). **d.** Heatmap (row-wise Z-scores) of the top 30 significantly reversed lipid species across individual biological replicates. **e.** Distribution of the top 10 lipid subclasses among the 109 significantly reversed lipid species. NC, normal control; IMQ, imiquimod; DEM, Demethylzeylasteral. In the panel labels, DEM denotes the IMQ+DEM group. Cer, ceramide; DG, diacylglycerol; FA, fatty acid; LPC, lysophosphatidylcholine; LPG, lysophosphatidylglycerol; MG, monoacylglycerol; PC, phosphatidylcholine; PE, phosphatidylethanolamine; PG, phosphatidylglycerol; PI, phosphatidylinositol; SM, sphingomyelin; SPH, sphingosine.
